# Overtone focusing in biphonic Tuvan throat singing

**DOI:** 10.1101/725267

**Authors:** Christopher Bergevin, Chandan Narayan, Joy Williams, Natasha Mhatre, Jennifer Steeves, Brad Story

## Abstract

Khoomei is a unique singing style originating from the Central Asian republic of Tuva. Singers produce two pitches simultaneously: a booming low-frequency rumble alongside a hovering high-pitched whistle-like tone. The biomechanics of this biphonation are not well-understood. Here, we use sound analysis, dynamic magnetic resonance imaging, and vocal tract modeling to demonstrate how biphonation is achieved by modulating vocal tract morphology. Tuvan singers show remarkable control in shaping their vocal tract to narrowly focus the harmonics (or overtones) emanating from their vocal cords. The biphonic sound is a combination of the fundamental pitch and a focused filter state, which is at the higher pitch (1-2 kHz) and formed by merging two formants, thereby greatly enhancing sound-production in a very narrow frequency range. Most importantly, we demonstrate that this biphonation is a phenomenon arising from linear filtering rather than a nonlinear source.

## Introduction

In the years preceding his death, Richard Feynman had been attempting to visit the small Asian republic of Tuva (***Leighton, 2000***). A key catalyst came from Kip Thorne, who had gifted him a record called *Melody tuvy*, featuring a Tuvan singing in a style known as “*Khoomei*”, or Xöömij. Although he was never successful in visiting Tuva, Feynman was nonetheless captivated by Khoomei, which can be best described as a high-pitched tone, similar to a whistle carrying a melody, hovering above a constant booming low-frequency rumble. Such is a form of biphonation, or in Feynman’s own words, “a man with two voices”. Khoomei, now a part of the UNESCO Intangible Cultural Heritage of Humanity, is characterized as “the simultaneous performance by one singer of a held pitch in the lower register and a melody … in the higher register” (***Aksenov, 1973***). How, indeed, does one singer produce two pitches at one time? Even today, the biophysical underpinnings of this biphonic human vocal style are not fully understood.

Normally, when a singer voices a song or speech, their vocal folds vibrate at a fundamental frequency (*f*_0_), generating oscillating airflow, forming the so-called *source*. This vibration however is not simply sinusoidal, as it also produces a series of harmonics tones (i.e., integer multiples of *f*_0_) (Fig.1). Harmonic frequencies in this sound above *f*_0_ are called overtones. Upon originating from the vocal folds, they are then sculpted by the vocal tract, which acts as a spectral *filter*. The vocal-tract filter has multiple resonances that accentuate certain clusters of overtones, creating *formants*. When speaking, we change the shape of our vocal tract to shift formants in systematic ways characteristic of vowel and consonant sounds. Indeed, singing largely uses vowel-like sounds (***Story et al., 2016***). However, in most singing, only one formant is emphasized (leading to a salient sense of dominant pitch) and the others contribute primarily to timbre. Khoomei has *two* strongly emphasized pitches, and the higher frequency formant can independently change and carry the melody of the song (***Kob, 2004***). Two possible loci for this biphonic property are the ‘source’ and/or the ‘filter’.

**Figure 1.**
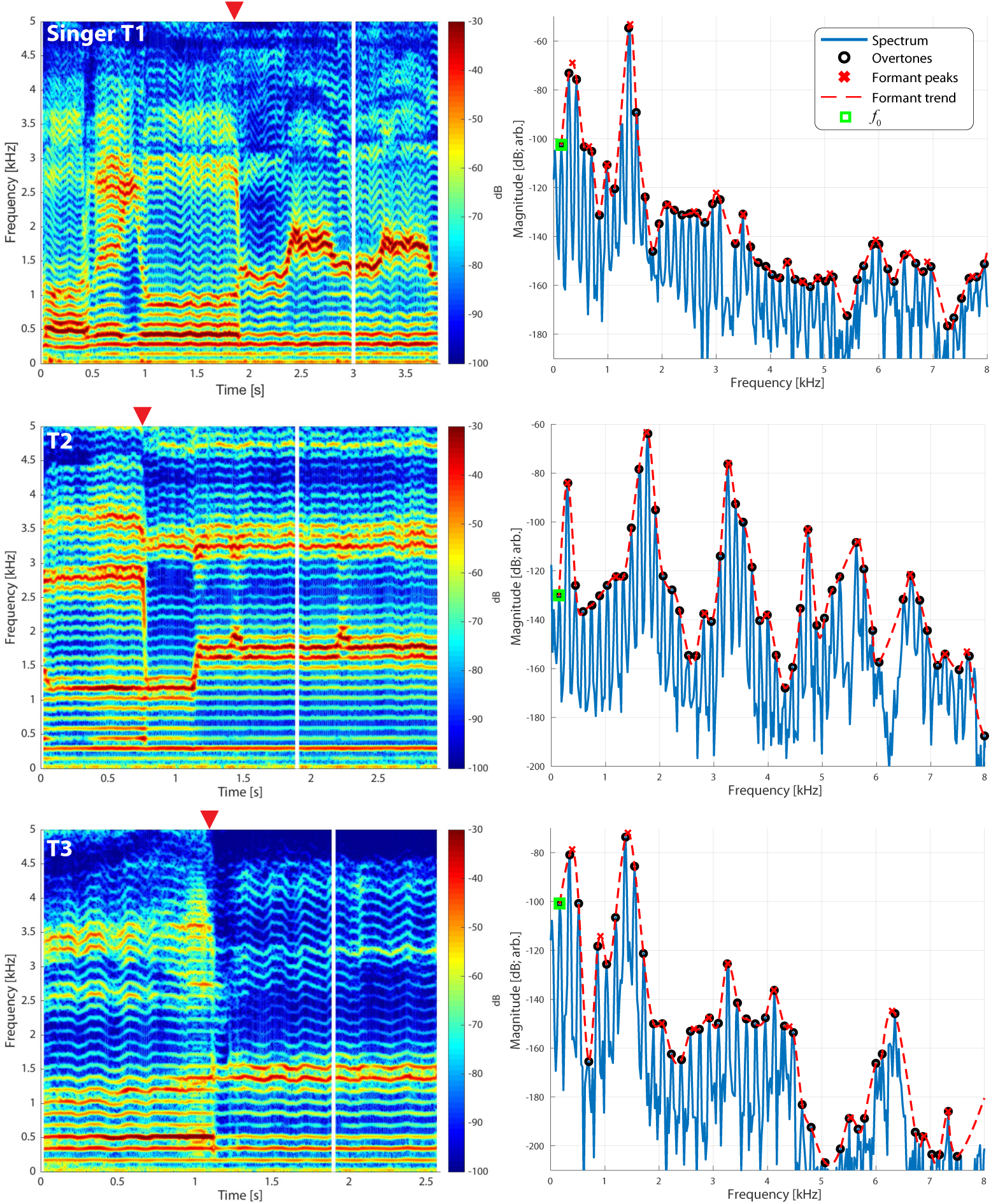
Frequency spectra of the songs three different singers transitioning from normal into biphonic singing. Vertical white lines indicate the time point for which the associated normal or biphonic song was extracted. Transition points from normal to biphonic singing state are denoted by the red triangle. (Right column). The fundamental frequency (*f*_0_) of the song which is indicated by a peak in the spectrum is marked by a green square. Overtones which represent integral multiples of this frequency are also indicated. Estimates of the formant structure are shown by overlaying a red stippled line and each formant peak is marked by an x. Note that the vertical scale is in decibels (e.g., a 120 dB difference is a million-fold difference in pressure amplitude). See also Figs. 4 and 2 for further quantification of these waveforms. The associated waveforms can be accessed in the Appendix [T1_3short.wav, T2_5short.wav, T3_2shortA.wav].

A source-based explanation could involve different mechanisms, such as two vibrating nonlinear sound sources in the syrinx of birds, which produce multiple notes that are harmonically unrelated (***Fee et al., 1998***; ***Zollinger et al., 2008***). Humans however are generally considered to have only a single source, the vocal cords. But there are an alternative possibilities: for instance, the source could be nonlinear and produce harmonically-unrelated sounds. For example, aerodynamic instabilities are known to produce biphonation (***Mahrt et al., 2016***). Further, Khoomei often involves dramatic and sudden transitions from simple tonal singing to biophonation (see Fig.1 and the *Appendix* for associated audio samples). Such abrupt changes are often considered hallmarks of physiological nonlinearity (***Goldberger et al., 2002***), and vocal production can generally be nonlinear in nature (***Herzel and Reuter, 1996***; ***Mergell and Herzel, 1997***; ***Fitch et al., 2002***; ***Suthers et al., 2006***). Therefore it remains possible that biphonation arises from a nonlinear source, i.e. the vocal cords.

Vocal tract shaping, a filter-based framework, provides an alternative explanation for biphonation. In one seminal study of Tuvan throat singing, Levin and Edgerton examined a wide variety of song types and suggested that there were three components at play. The first two (“tuning a harmonic” relative to the filter and lengthening the closed phase of the vocal fold vibration) represented a coupling between source and filter. But it was the third, narrowing of the formant, that appeared crucial. Yet, the authors offered little empirical justification for how these effects are produced by the vocal tract shape in the presented radiographs. Thus it remains unclear how the high-pitched formant in Khoomei was formed (***Grawunder, 2009***). Another study (***Adachi and Yamada, 1999***) examined a throat singer using magnetic resonance imaging (MRI) and captured static images of the vocal tract shape during singing. These images were then used in a computational model to produce synthesized song. Adachi and Yamada argued that a “rear cavity” was formed in the vocal tract and its resonance was essential to biphonation. However, their MRI data reveal limited detail since they were static images of singers already in the biphonation state. Small variations in vocal tract geometry can have pronounced effects on produced song (***Story et al., 1996***). Data from static MRI would reveal little about how and which parts of the vocal tract change shape as the singers transition from simple tonal song to biphonation. To understand which features of vocal tract morphology are crucial to biophonation, a dynamic description of vocal tract morphology would be required.

Here we study the dynamic changes in the vocal tracts of multiple expert practitioners from Tuva as they produce Khoomei. We use MRI to acquire volumetric 3D shape of the vocal tract of a singer during biphonation. Then, we capture the dynamic changes in a midsagittal slice of the vocal tract as singers transition from tonal to biphonic singing while making simultaneous audio recordings of the song. We use these empirical data to guide our use of a computational model, which allows us to gain insight into which features of vocal tract morphology are responsible for the singing phonetics observed during biophonic Khoomei song [e.g., ***Story et al. (2016)***]. We focus specifically on the Sygyt (or Sigit) style of Khoomei (***Aksenov, 1973***).

## Results

### Audio recordings

We made measurements from three Tuvan singers performing Khoomei in the Sygyt style (designated as T1–T3) and one (T4) in a non-Sygyt style. Songs were analyzed using short-time Fourier transforms (STFT), which provide detailed information in both temporal and spectral domains. We recorded the singers transitioning from normal singing into biphonation, Fig.1 showing this transition for three singers. The *f*_0_ of their song is marked in the figure (approximately 140 Hz for subject T2, 164 Hz for both T1 and T3) and the overtone structure appears as horizontal bands. Varying degrees of vibrato can be observed, dependent upon the singer (Fig.1; see also longer spectrograms in Figs. 6 and 7). Most of the energy in their song is concentrated in the overtones and no subharmonics (i.e., peaks at half-integer multiples of *f*_0_) are observed. In contrast to these three singers, singer T4 performing in a non-Sygyt style exhibited a fundamental frequency of approximately 130 Hz, although significant energy additionally appears around 50-55 Hz, well below an expected subharmonic (Fig.5).

**Figure 2.**
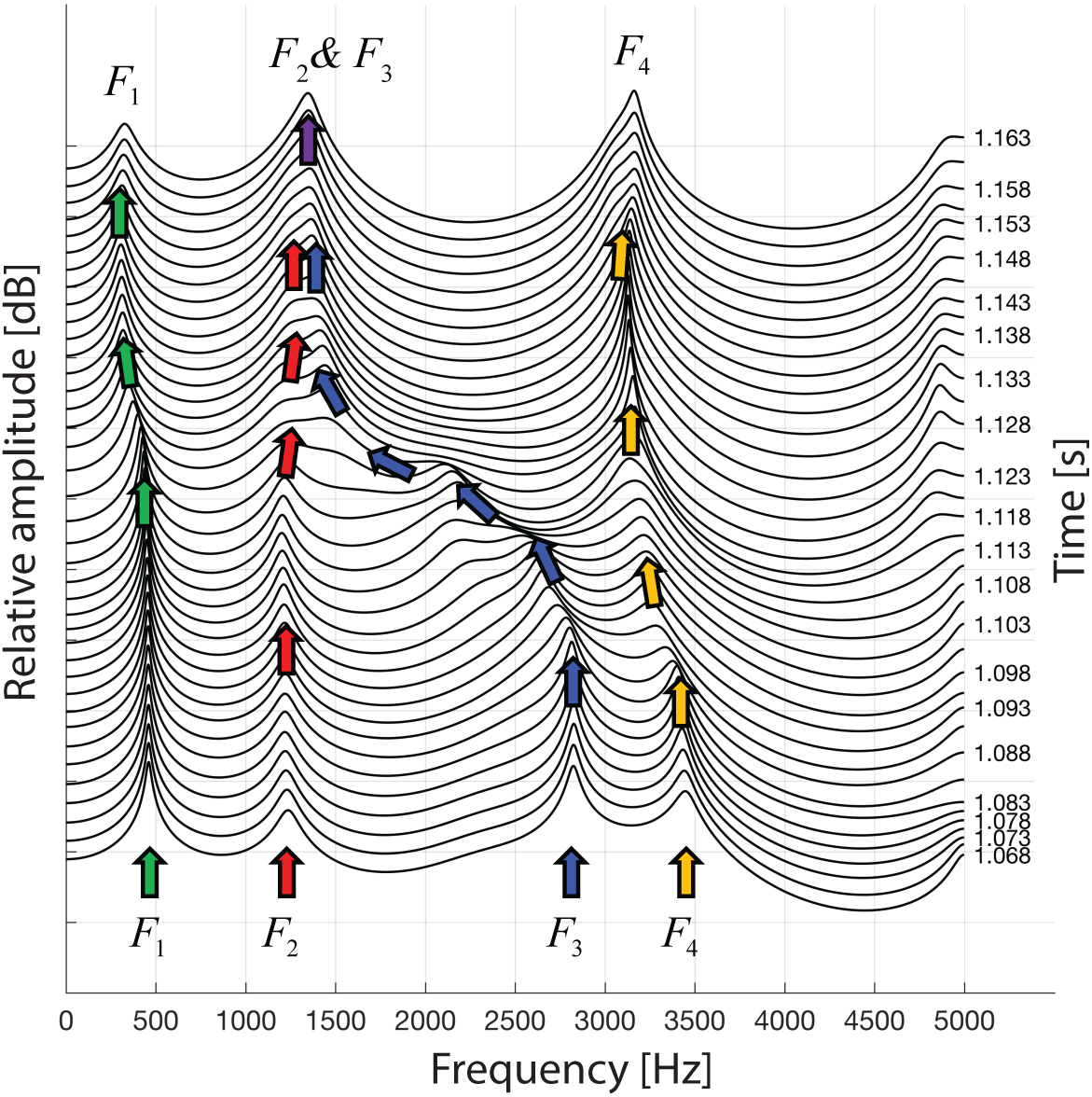
A waterfall plot representing the spectra at different time points as singer T2 transitions from normal singing into biphonation (T2_3short.wav). The superimposed arrows are color-coded to help visualize how the formants change about the transition, chiefly with F3 shifting to merge with F2. This plot also indicates the second focused state centered just above 3 kHz is a sharpened F4 formant.

**Figure 3.**
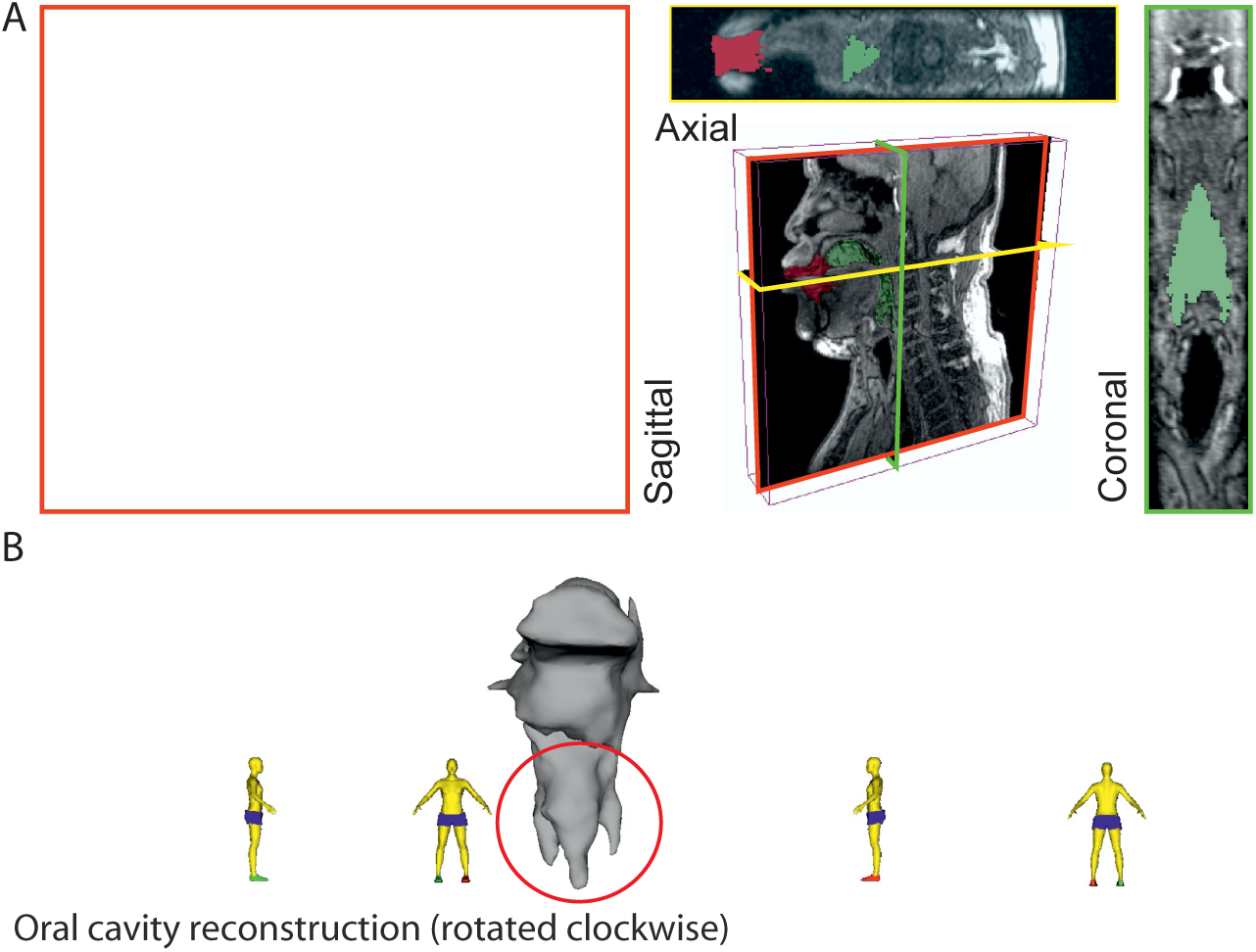
3-D reconstruction of volumetric MRI data taken from singer T2 (Run3; see *Appendix*, including Fig.19). (A) Example of MRI data sliced through three different planes, including a pseudo-3D plot. Airspaces were determined manually (green areas behind tongue tip, red for beyond). Basic labels are included: L – lips, J – jaw, T– tongue, AR – aveolar ridge, V – velum, E – epiglottis, Lx – larynx, and T – trachea. The shadow from the dental post is visible in the axial view on the left hand side and stops near the midline leaving that view relatively unaffected. (B) Reconstructed airspace of the vocal tract from four different perspectives. The red circle highlights the presence of the piriform sinuses (***Dang and Honda, 1997***).

**Figure 4.**
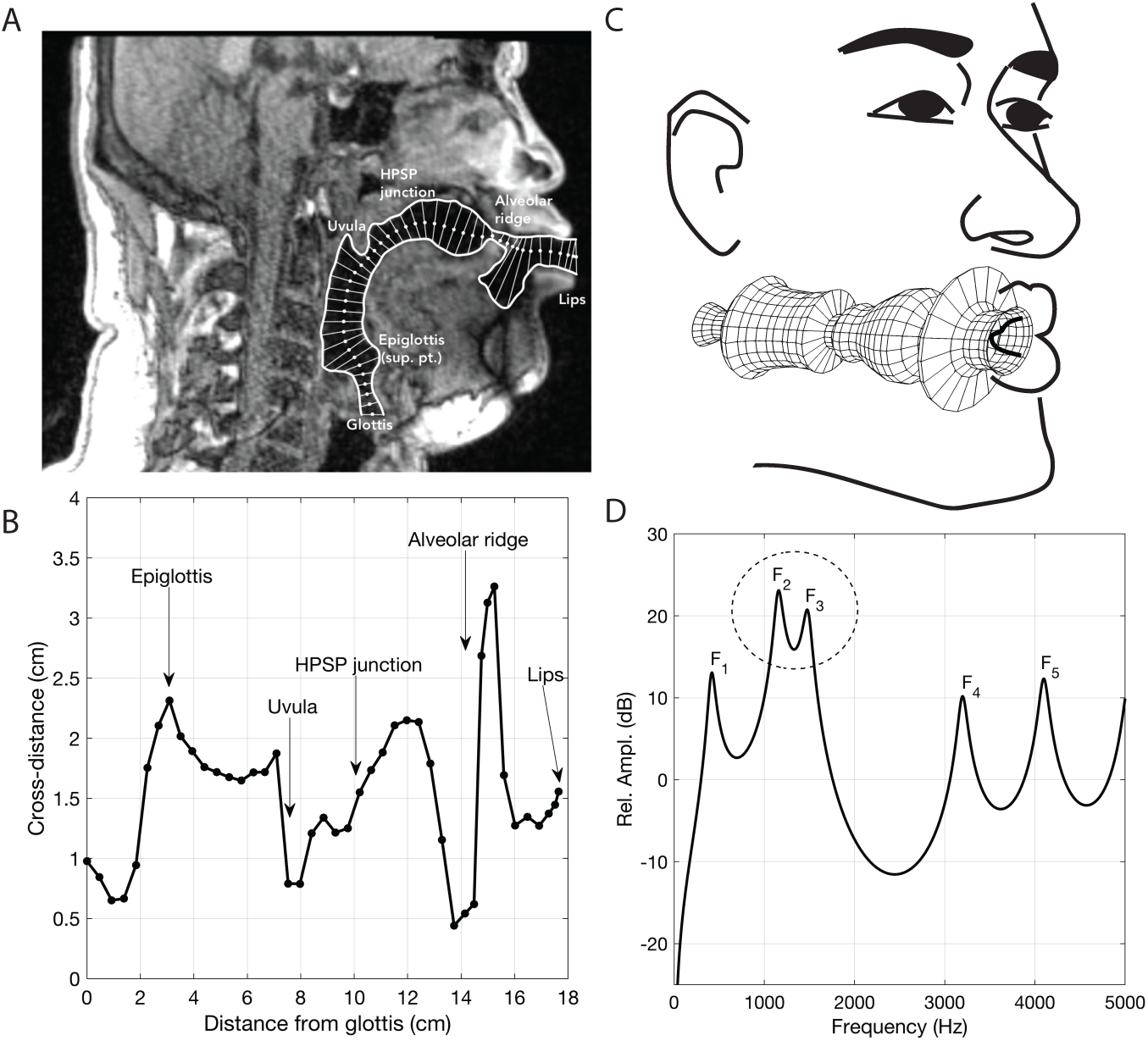
Analysis of vocal tract configuration during singing. (A) 2D measurement of tract shape. The inner and outer profiles were manually traced, whereas the centerline (white dots) was found with an iterative bisection technique. The distance from the inner to outer profile was measured along a line perpendicular to each point on the centerline (thin white lines). (B) Collection of cross-distance measurements plotted as a function of distance from the glottis. Area function can be computed directly from these values and is derived by assuming the cross-distances to be equivalent diameters of circular cross-sections (see *Methods*). (C) Schematic indicating associated modeling assumptions, including vocal tract configuration as in panel B [adapted from ***Bunton et al. (2013)***, under a Creative Commons CC-BY licence, https://creativecommons.org/licenses/by/4.0/]. (D) Model frequency response calculated from the associated area function stemming from panels B & C. Each labeled peak can be considered a formant frequency and the dashed circle indicates merging of formants F2 and F3.

**Figure 5.**
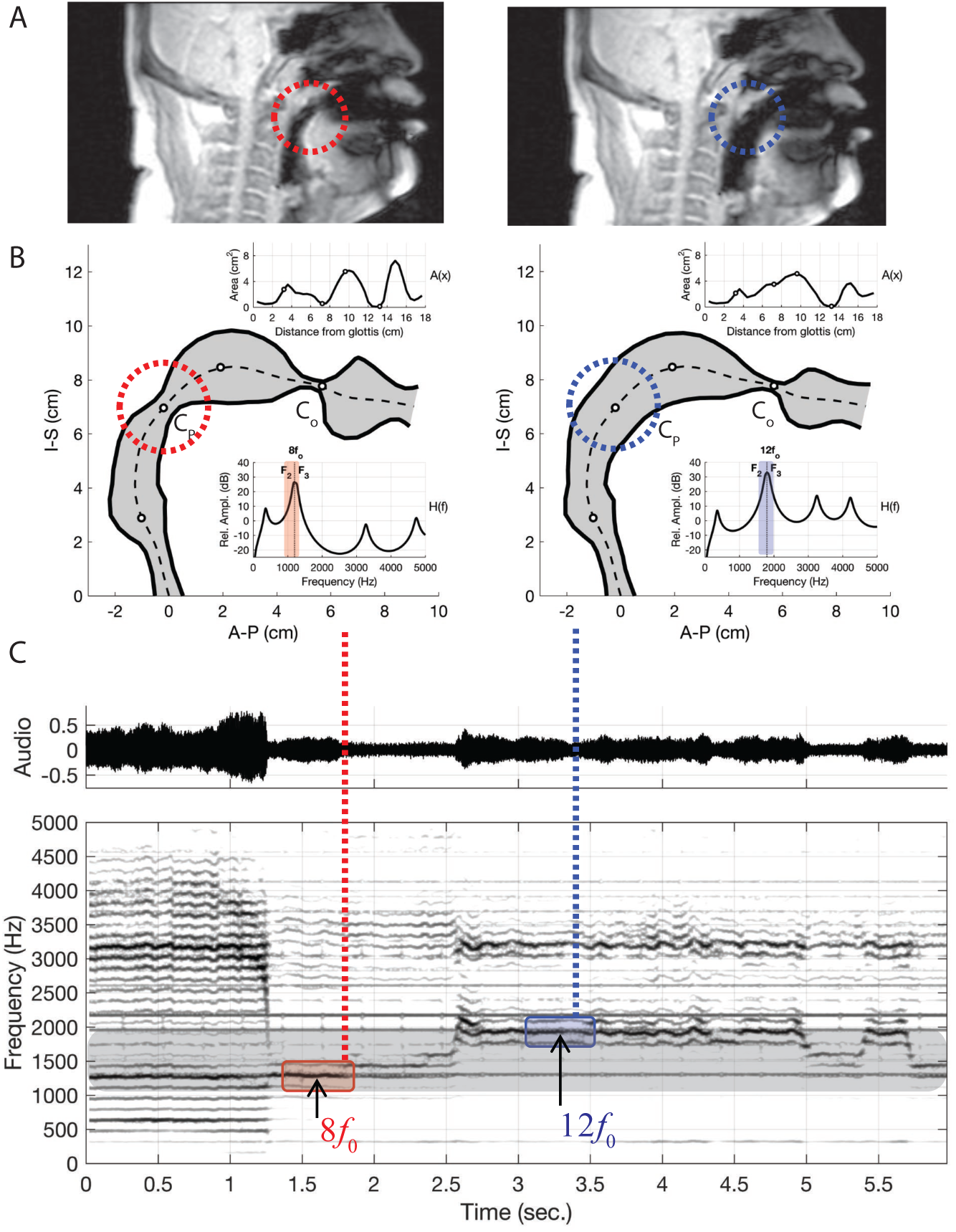
Results of changing vocal tract morphology in the model by perturbing the baseline area function *A*_0_(*x*) to demonstrate the merging of formants *F*_2_ and *F*_3_, atop two separate overtones as apparent in the two columns of panels A and B. (A) The frames from dynamic MRI with red and blue dashed circles highlighting the location of the key vocal tract constrictions. (B) Model-based vocal tract shapes stemming from the MRI data, including both the associated area functions (top inset) and frequency response functions (bottom inset). *C*_*o*_ indicates the constriction near the alveolar ridge while *C*_*P*_ the constriction near the uvula in the upper pharynx. (C) Waveform and corresponding spectrogram of audio from singer T2 (a spectrogram from the model is shown in *Appendix* Fig.14). Note that the merged formants lie atop either the 7th overtone (i.e., 8*f*_0_) or the 11th ((i.e., 12*f*_0_).

If we take a slice, i.e. a time-point from the spectrogram and plot the spectrum, we can observe the peaks to infer the formant structure from this representation of the sound (red-dashed lines in Fig.1 and Appendix Fig.4). As the singers transition from normal singing to biphonation, we see that the formant structure changes significantly and the positions of formant peaks shift dramatically and rapidly. Note that considering time points before and after the transitions also provides an internal control for both normal and focused song types (Fig. 4). Once in the biphonation mode, all three singers demonstrate overtones in a narrow spectral band around 1.5-2 kHz; we refer to this as the *focused state*. Specifically, Fig.1 shows that not only is just a single or small group of overtones accentuated, but also that nearby ones are greatly attenuated: ±1 overtones are as much 15–35 dB and ±2 overtones are 35–65 dB below the central overtone. Additionally, in the focused state, the fundamental frequency is also relatively small. In normal singing, prior to the focused state, the majority of vocal energy is below 1 kHz and the *f*_0_ is strongly voiced, whereas after the transition the strongly voiced component of the song shifts to whistle-like pitch in the 1.5-2 kHz region.

To objectively and quantitatively assess the degree of focus, we computed an energy ratio *e*_*R*_(*f*_*L*_, *f*_*H*_) that characterizes the relative degree of energy brought into a narrow band against the energy spread over the full spectrum occupied by human speech (see *Methods*). In normal speech and singing, for [*f*_*L*_, *f*_*H*_] = [1, 2 kHz], typically *e*_*R*_ is small (i.e., energy is spread across the spectrum, not *focused* into that narrow region between 1 and 2 kHz). For the Tuvan singers, prior to a transition into a focused state, *e*_*R*_(1, 2) is similarly small. However once the transition occurs (red triangle in Fig.1), those values are large (upwards of 0.5 and higher) and sustained across time (*Appendix* Figs. 2 & 3). For one of the singers (T2) the situation was more complex, as he created multiple focused formants (Fig.1 middle panels and *Appendix* Figs. 6,8). The second focused state was not explicitly dependent upon the first: The first focused state clearly moves and transitions between approximately 1.5–2 kHz (by 30%) while the second focused state remains constant at approximately 3–3.5kHz (changing less than 1%). Thus the focused states are not harmonically related. Unlike the other singers, T2 not only has a second focused state, but also had more energy in the higher overtones (Fig.1). As such, singer T2 also exhibited a different *e*_*R*_ timecourse, which took on values that could be relatively large even prior to the transition. This may be because he took multiple ways to approach the transition into a focused state (e.g., *Appendix* Fig.9).

Plotting spectra around the transition from normal to biphonation singing in a waterfall plot indicates that the sharp focused filter is achieved by merging two broader formants together (*F*_2_ and *F*_3_ in Fig.2) (***Kob, 2004***). This transition into the focused state is fast (∼40–60 ms), as are the shorter transitions within the focused state where the singer melodically changes the filter that forms the whistle-like component of their song [Fig.1, *Appendix* Fig.8].

### Vocal tract MRI

While we can infer the shape of the formants in Khoomei by examining audio recordings, such analysis is not conclusive in explaining the mechanism used to achieve these formants. The working hypothesis is that vocal tract shape determines these formants, therefore it is crucial to examine the shape and dynamics of the vocal tract to determine whether the acoustic measurements are consistent with this hypothesis. In order to accomplish this, we obtained MRI data from one of the singers (T2) that are unique in two regards. First, there are two types of MRI data reported here: steady-state volumetric data (Fig.3 and *Appendix* Fig.19) and dynamic midsagittal images at several frames per second that capture changes in vocal tract position (Fig.4A–B and *Appendix* Fig.21). Second is that the dynamic data allow us to examine vocal tract changes as song transitions into a focused state (e.g., *Appendix* Fig.21).

The human vocal tract begins at the vocal folds and ends at the lips. Airflow produced by the vocal cords sets the air-column in the tract into vibration, and its acoustics determine the sound that emanates from the mouth. The vocal tract is effectively a tube-like cavity whose shape can be altered by several articulators: the jaw, lips, tongue, velum, epiglottis, larynx and trachea (Fig.3). Producing speech or song requires that the shape of the vocal tract, and hence its acoustics, are precisely controlled (***Story et al., 2016***).

Several salient aspects of the vocal tract during the production of Khoomei can be observed in the volumetric MRI data. The most important feature however, is that there are two distinct and relevant constrictions when in the focused state, corresponding roughly to the uvula and alveolar ridge. Additionally, the vocal tract is expanded in the region just anterior to the alveolar ridge (Fig.3A). The retroflex position of the tongue tip and blade produces a constriction at 14 cm, and also results in the opening of this sublingual space. It is the degree of constriction at these two locations that is hypothesized to be the primary mechanism for creating and controlling the frequency at which the formant is *focused*.

### Modeling

Having established that the shape of vocal tract during Khoomei does indeed have two constrictions, as observed by other groups, the primary goals of our vocal tract modeling efforts were to use the dynamic MRI data as morphological benchmarks and capture the merging of formants to create the focused states as well as the dynamic transitions into them. Our approach was to use a well-established linear “source/filter” model [e.g., ***Stevens (2000***)] that includes known energy losses (***Sondhi and Schroeter, 1987***; ***Story et al., 2000***; ***Story, 2013***). Here, the vibrating vocals folds act as the broadband sound source (with the *f*_0_ and associated overtone cascade), while resonances of the vocal tract, considered as a series of 1-D concatenated tubes of variable uniform radius, act as a primary filter. We begin with a first order assumption that the system behaves linearly, which allows us for a simple multiplicative relationship between the source and filter in the spectral domain (e.g., *Appendix* Fig.10).

Acoustic characteristics of the vocal tract can be captured by transforming the three-dimensional configuration (Fig. 3) into a tube with variation in its cross-sectional area from the glottis to the lips (Figs. 4 and 5). This representation of the vocal tract shape is called an *area function*, and allows for calculation of the corresponding frequency response function (from which the formant frequencies can be determined) with a one-dimensional wave propagation algorithm. Although the area function can be obtained directly from a 3D vocal tract reconstruction [e.g., ***Story et al.*** (***1996***)], the 3D reconstructions of the Tuvan singer’s vocal tract were affected by a large shadow from a dental post (Fig. 2) and were not amenable to detailed measurements of cross-sectional area. Instead, a cross-sectional area function was measured from the midsagittal slice of the 3D image set (see *Methods* and *Appendix* for details). Thus, the MRI data provided crucial bounds for model parameters: the locations of primary constrictions and thereby the associated area functions.

The frequency response functions derived from the above static volumetric MRI data (e.g., Fig.4D) indicate that two formants *F*_2_ and *F*_3_ cluster together, thus enhancing both their amplitudes. Clearly, if *F*_2_ and *F*_3_ could be driven closer together in frequency, they would merge and form a single formant with unusually high amplitude. We hypothesize that this mechanism could be useful for effectively amplifying a specific overtone, such that it becomes a prominent acoustic feature in the sound produced by a singer, specifically the high frequency component of Khoomei.

Next, we used the model in conjunction with time-resolved MRI data to investigate how the degree of constriction and expansion at different locations along the vocal tract axis could be a mechanism for controlling the transition from normal to overtone singing and the pitch while in the focused state. These results are summarized in Fig.4 (further details are in the *Appendix*). While the singers are in the normal song mode, there are no obvious strong constrictions in their vocal tracts (e.g., *Appendix* Fig.11). After they transition, in each MRI from the focused state, we observe a strong constriction near the aveolar ridge. We also observe a constriction near the uvula in the upper pharynx, but the degree of constriction here varies. If we examine the simultaneous audio recordings, we find that variations in this constriction are co-variant with the frequency of the focused formant. From this, we surmise that the mechanism for controlling the enhancement of voice harmonics is the degree of constriction near the alveolar ridge in the oral cavity (labeled *C*_*o*_ in Fig. 4), which affects the proximity of *F*_2_ and *F*_3_ to each other (*Appendix* Fig.12). Additionally, the degree of constriction near the uvula in the upper pharynx (*C*_*P*_) controls the actual frequency at which *F*_2_ and *F*_3_ converge (*Appendix* Fig.13). Other parts of the vocal tract, specifically the expansion anterior to *C*_*o*_, may also contribute since they also show small co-variations with the focused formant frequency (*Appendix* Fig.14). Further, a dynamic implementation of the model, as shown in *Appendix* Fig.14, reasonably captures the rapid transition into/out of the focused state as shown in Fig.1. Taken together, the model confirms and explains how these articulatory changes give rise to the observed acoustic effects.

To summarize, an overtone singer could potentially “play” (i.e., select) various harmonics of the voice source by first generating a tight constriction in the oral cavity near the alveolar ridge, and then modulating the degree of constriction in the uvular region of the upper pharynx to vary the position of the focused formant, thereby generating a basis for melodic structure.

## Discussion

This study has shown that Tuvan singers performing Sygyt-style Khoomei exercise precise control of the vocal tract to effectively merge multiple formants together. They morph their vocal tracts so to create a sustained *focused* state that effectively filters an underlying stable array of overtones. This focused filter greatly accentuates energy of a small subset of higher order overtones primarily in the octave-band spanning 1–2 kHz, as quantified by an energy ratio *e*_*R*_(1, 2). Some singers are even capable of producing additional foci at higher frequencies. Below, we argue that a linear framework [i.e., source/filter model, (***Stevens, 2000***)] appears sufficient to capture this behavior including the sudden transitions into a focused state, demonstrating that nonlinearities are not *a priori* essential. That is, since the filter characteristics are highly sensitive to vocal tract geometry, precise biomechanical motor control of the singers is sufficient to achieve a focused state without invoking nonlinearities found in other vocalization types [e.g., ***Fee et al.*** (***1998***)]. Additionally, we suggest the possibility that a unique feature of the cochlea may affect how focused overtone song is encoded, thereby enhancing the saliency of the perception of this type of vocalization.

### Source or filter?

The notion of a focused state is mostly consistent with vocal tract filter-based explanations for biphonation in previous studies [e.g., ***Bloothooft et al. (1992)***; ***Edgerton et al. (1999)***; ***Adachi and Yamada (1999)***; ***Grawunder (2009)***], where terms such as an “interaction of closely spaced formants”, “reinforced harmonics”, and “formant melting” were used. Additionally, the merging of multiple formants is closely related to the “singer’s formant”, which is proposed to arise around 3 kHz due to formants *F*_3_–*F*_5_ combining (***Story et al., 2016)***, though this is typically more broad and less prominent than the focused states exhibited by the Tuvans. Our results explain how this occurs and are also broadly consistent with Adachi & Yamada (***Adachi and Yamada, 1999)*** in that a constricted “rear cavity” is crucial. However, we find that this rear constriction determines the pitch of the focused formant, whereas it is the “front cavity” constriction near the alveolar ridge that produces the focusing effect (i.e., merging of formants *F*_2_ and *F*_3_).

Further, the present data appear in several ways inconsistent with conclusions from previous studies of Khoomei, especially those that center on effects that arise from changes in the source. Three salient examples are highlighted. First, we observed overtone structure to be highly stable, though some vibrato may be present. This contrasts the claim by Levin and Edgerton that “(t)o tune a harmonic, the vocalist adjusts the fundamental frequency of the buzzing sound produced by the vocal folds, so as to bring the harmonic into alignment with a formant” (***Levin and Edgerton, 1999)***. That is, we see no evidence for the overtone “ladder” being lowered or lifted as they suggested (note in Fig.1, *f*_0_ is held nearly constant). Second, a single sharply defined harmonic alone is not sufficient to get the salient perception of a focused state, as had been suggested by ***Levin and Edgerton (1999)***. Consider *Appendix* Fig.9, especially at the 4 s mark, when “pressed”^1^ transition (***Lindestad et al., 2001***; ***Edmondson and Esling, 2006)*** phonation occurs prior to onset perceptually of the focused state. There, a harmonic at 1.51 kHz dominates (i.e., the two flanking overtones are approximately 40 dB down), yet the song has not yet perceptibly transitioned. It is not until the cluster of overtones at 3–3.5 kHz is brought into focus that the perceptual effect becomes salient, perhaps because prior to the 5.3 s mark the broadband nature of those frequencies effectively masks the first focused state. Third, we do not observe subharmonics, which contrasts a prior claim (***Lindestad et al., 2001)*** that “(t)his combined voice source produces a very dense spectrum of overtones suitable for overtone enhancement”. However, that study was focused on a different style of song called “Kargyraa”, which does not exhibit as clearly a focused state as in Sygyt.

### Linear versus nonlinear mechanisms

An underlying biophysical question is whether focused overtone song arises from inherently linear or nonlinear processes. Given that Khoomei consists of the voicing of two or more pitches at once and exhibits dramatic and fast transitions from normal singing to biphonation, nonlinear phenomena may seem like an obvious candidate (***Herzel and Reuter, 1996)***^2^. For example, certain frog species exhibit biphonation, and it has been suggested that their vocalizations can arise from complex nonlinear oscillatory regimes of separate elastically coupled masses (***Suthers et al., 2006)***. Second is sudden transition into the focused state, as seen in Fig.1. The appearance of abrupt changes in physiological systems has been argued to be a flag for nonlinear mechanisms (***Goldberger et al., 2002)***, such as by virtue of progression through a bifurcation.

Our results present two lines of evidence that argue against Sygyt-style Khoomei arising primarily from a nonlinear process. First, the underlying harmonic structure of the vocal fold source appears highly stable through the transition into the focused state (Fig.1). There is little evidence of subharmonics. A source spectral structure that is comprised of an *f*_0_ and integral harmonics would suggest a primarily linear source mechanism. Second is that our modeling efforts, which are chiefly linear in nature, reasonably account for the sudden and salient transition. That is, the model is readily sufficient to capture the characteristic that small changes in the vocal tract can produce large changes in the filter. Thereby, precise and fast motor control of the articulators in a linear framework accounts for the transitions into and out of the focused state. Thus in essence, Sygyt-style Khoomei could be considered a linear means to achieve biphonation. Connecting back to nonlinear phonation mechanisms in non-mammals, our results provide further context for how human song production and perception may be similar and/or different relative to non-humans [e.g., ***Doolittle et al. (2014)***; ***Kingsley et al. (2018)***].

Features that appear transiently in spectrograms do provide hints of source nonlinearity though, such as the brief appearance of subharmonics in some instances (*Appendix* Fig.15B). Such provides an opportunity to address limitations of the current modeling efforts and highlight future considerations. We suggest that further analysis [e.g., ***Theiler et al. (1992)***; ***Tokuda et al. (2002)***; ***Kantz and Schreiber (2004)***] on Khoomei audio recordings may help inform the model and better capture focused filter sharpness and the origin of secondary focused states. Several potential areas for improvement are: nonlinear source–filter coupling (***Titze, 2008)***, a detailed model of glottal dynamics [e.g., ratio of open/closed phases in glottal flow ***Grawunder (2009)***; ***Li and Hou (2017)***, periodic vibrations in *f*_0_], inclusion of piriform sinuses as side-branch resonators (***Dang and Honda, 1997***; ***Titze and Story, 1997)***, inclusion of the 3-D geometry (Fig.3), and detailed study of the front cavity (e.g., lip movements) that may be used by the singer to maintain control of focused state as well as make subtle manipulations.

### Exploiting a cochlear transition?

A previous study considered perception of focused overtone song and reported several psychoa-coustic measurements (***Bloothooft et al., 1992)***. They noted listeners placed overtone sounds in a different category from vowels sung more conventionally, suggesting this dichotomy stems from an independent perception of the frequency of the overtone (***Bloothooft et al., 1992)***. Is there something different about the way focused overtone song is perceived given the human auditory pathway? Recent advancements suggest that this may in fact be the case by virtue of a *transition* in the cochlea.

In terms of auditory transduction, several considerations motivate the notion of “two cochleas” (***Shera, 2015)***. That is, there is increasing evidence that suggests auditory information is processed differently between the base and apex of the cochlea, with a transition occurring around 1–1.5 kHz (***Shera et al., 2010)***. First, while the tonotopic map of the cochlea is roughly exponential (i.e., one octave equates to a fixed distance), this relationship deviates at lower frequencies below ∼1 kHz [e.g., ***Greenwood (1990)***; see *Appendix* Fig.18]. Second, scaling symmetry, a key principle commonly evoked in modeling cochlear mechanics, appears to break down below 1–1.5 kHz (***van der Heijden and Joris, 2006***; ***Abdala et al., 2011)***. Third, various neurophysiological measures [e.g., ***van der Heijden and Joris (2006)***; ***Verschooten et al. (2018)***] suggest auditory nerve responses encode information differently between base and apex, with a steep drop in the ability to encode timing information above 1–2 kHz (***Verschooten et al., 2018)***. Fourth, it is well known that harmonic components of perceived pitch transition from resolved to unresolved around 1–2 kHz (***Plack et al., 2006)***. Lastly, advances in cochlear imaging show that the micromechanical vibrations of the cochlea are different between base and apex [e.g., ***Dong et al. (2018)***], with large differences in tuning sharpness. While recent modeling efforts have made progress in reconciling these differences in apical and basal sound processing for mammals (***Sasmal and Grosh, 2019)***, it has been suggested that the human cochlea may be different in many regards from the canonical viewpoint of a relatively generic cochlear structure/physiology/function across mammals (***Joris et al., 2011***; ***Raufer et al., 2019)***. Typically, adult phonetic signals contain significant energy below 1 kHz (***Stevens, 2000)*** and this transition thereby may not strongly affect speech perception. However focused overtone song concentrates energy narrowly into the 1–2 kHz octave band, and lowers energy below 1 kHz. Taken together, these lines of evidence suggest that Sygyt-style Khoomei may elicit such a salient percept in part because that phonetic overtone energy of higher order harmonics (e.g., 8–12 *f*_0_) is focused to straddle the region between those two functionally distinct parts of the human cochlea.

## Methods

### Acoustical recordings

Recordings were made at York University (Toronto, ON, Canada) in a double-walled acoustic isolation booth (IAC) using a Zoom H5 24-bit digital recorder and an Audio-Technica P48 condenser microphone. A sample rate of 96 kHz was used. Spectral analysis was done using custom-coded software in Matlab. Spectrograms were typically computed using 4096 point window segments with 95% fractional overlap and a Hamming window. Harmonics (black circles in Fig.1) were estimated using a custom-coded peak-picking algorithm. Estimated formant trends (red dashed lines in Fig.1) were determined via a lowpass interpolating filter built into Matlab’s digital signal processing toolbox with a scaling factor of 10. From this trend, the peak-picking was reapplied to determine “formant” frequencies (red x’s in Fig.1). This process could be repeated across the spectrogram to effectively track overtone and formant frequency/strength, as shown in Fig.1.

To quantify the focused states, we developed a dimension-less measure *e*_*R*_(*f*_*L*_, *f*_*H*_) to represent the energy ratio of that spanning a frequency range *f*_*H*_ - *f*_*L*_ relative to the entire spectral output. This can be readily computed from the spectrogram data as follows. First take a “slice” from the spectrogram and convert spectral magnitude to linear ordinate and square it (as intensity is proportional to pressure squared). Then integrate across frequency, first for a limited range spanning [*f*_*L*_, *f*_*H*_] (e.g., 1–2 kHz) and then a broader range of [0, *f*_*max*_] (e.g., 0–8 kHz; 8 kHz is a suitable maximum as there is little acoustic energy in vocal output above). The ratio of these two is then defined as *e*_*R*_, and takes on values between 0 and 1. This can be expressed more explicitly as:

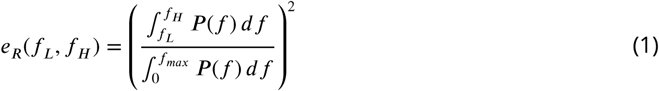

where *P* is the magnitude of the scaled sound pressure, *f* is frequency, and *f*_*L*_ & *f*_*H*_ are filter limits for considering the focused state. The choice of [*f*_*L*_, *f*_*H*_] = [1, 2] kHz has the virtue of spanning an octave, which also closely approximates the “seventh octave” from about C6 to C7. *e*_*R*_ did not depend significantly upon the length of FFT window. Values of *e*_*R*_ for the waveforms used in Fig.1 are shown in Figs.2 and 3.

### MRI Acquisition & Volumetric Analysis

MRI images were acquired at the York MRI Facility on a 3.0 Tesla MRI scanner (Siemens Magnetom TIM Trio, Erlangen, Germany), using a 12-channel head coil and a neck array. Data were collected with approval of the York University Institutional Review Board. The participant was fitted with an MRI compatible noise cancelling microphone (Optoacoustics, Mazor, Israel) mounted directly above the lips. The latency of the microphone and noise cancelling algorithm was 24 ms. Auditory recordings were made in QuickTime on an iMac during the scans to verify performance. Images were acquired using one of two paradigms, static or dynamic. Static images were acquired using a T1-weighted 3D gradient echo sequence in the sagittal orientation with 44 slices centered on the vocal tract, TR = 2.35 ms, TE = 0.97 ms, flip angle = 8 degrees, FoV = 300 mm, and a voxel dimension of 1.2 × 1.2 × 1.2 mm. Total acquisition time was 11 seconds. The participant was instructed to begin singing a tone, and to hold it in a steady state for the duration of the scan. The scan was started immediately after the participant began to sing and had reached a steady state. Audio recordings verified a consistent tone for the duration of the scan. Dynamic images were acquired using a 2D gradient echo sequence. A single 10.0 mm thick slice was positioned in a sagittal orientation along the midline of the vocal tract, TR = 4.6 ms, TE = 2.04 ms, flip angle = 8 degrees, FoV = 250 mm, and a voxel dimension of 2.0 × 2.0 × 10.0 mm. One hundred measurements were taken for a scan duration of 27.75 seconds. The effective frame rate of the dynamic images was 3.6 Hz. Audio recordings were started just prior to scanning. Only subject T2 participated in the MRI recordings. The participant was instructed to sing a melody for the duration of the scan, and took breaths as needed.

For segmentation (Fig.3), 3D MRI Images (Run1; see *Appendix*) were loaded into Slicer (version 4.6.2 r25516). The air-space in the oral cavity was manually segmented using the segmentation module, identified and painted in slice by slice. Careful attention was paid to the parts of the oral cavity that were affected by the artifact from the dental implant. The air cavity was manually repainted to be approximately symmetric in this region using the coronal and axial view (Fig.3A). Once completely segmented, the sections were converted into a 3D model and exported as a STL file. This mesh file was imported into MeshLab (v1.3.4Beta) for cleaning and repairing the mesh. The surface of the STL was converted to be continuous by removing non-manifold faces and then smoothed using depth and Laplacian filters. The mesh was then imported into Meshmixer where further artifacts were removed. This surface smoothed STL file was finally reimported into Slicer generating the display in Fig.3B.

### Computational modeling

Measurement of the cross-distance function is illustrated in Fig. 4. The inner and outer profiles of the vocal tract were first determined by manual tracing of the midsagittal image. A 2D iterative bisection algorithm (***Story, 2007)*** was then used to find the centerline within the profiles extending from the glottis to the lips, as shown by the white dots in Fig. 4A. Perpendicular to each point on the centerline, the distance from the inner to outer profiles was measured to generate the cross-distance function shown in Fig. 4B; the corresponding locations of the anatomic landmarks shown in the midsagittal image are also indicated on the cross-distance function.

The cross-distance function, *D*(*x*), can be transformed to an approximate area function, *A*(*x*), with the relation *A*(*x*) = *kD*^*α*^ (*x*), where *k* and *α* are a scaling factor and exponent, respectively. If the elements of *D*(*x*) are considered to be diameters of a circular cross-section, *k* = (*π*/4) and *α* = 2. Although other values of *k* and *α* have been proposed to account for the complex shape of the vocal tract cross-section (***Heinz and Stevens, 1964***; ***Lindblom and Sundberg, 1971***; ***Mermelstein, 1973)***, there is no agreement on a fixed set of numbers for each parameter. Hence, the circular approximation was used in this study to generate an estimate of the area function. In panel of Fig. 4C, the area function is plotted as its tubular equivalent where the radii *D*(*x*)/2 were rotated about an axis to generate circular sections from glottis to lips.

The associated frequency response of that area function is shown in Fig. 4D and was calculated with a transmission line approach (***Sondhi and Schroeter, 1987***; ***Story et al., 2000)*** that included energy losses due to yielding walls, viscosity, heat conduction, and acoustic radiation at the lips; side branches such the piriform sinuses were not considered in detail for this study. The first five formant frequencies (resonances), *F*_1_, …, *F*_5_, were determined by finding the peaks in the frequency response functions with a peak-picking algorithm(***Titze et al., 1987)*** and are located at 400, 1065, 1314, 3286, and 4029 Hz, respectively.

To examine changes in pitch, a particular vocal tract configuration was manually “designed” (Fig. 11a) such that it included constrictive and expansive regions at locations similar to those measured from the singer (i.e., Fig. 4), but to a less extreme degree. We henceforth denote this area function as *A*_0_(*x*), and it generates a frequency response with widely spaced formant frequencies (*F*_1…5_ = [529, 1544, 2438, 3094, 4236] Hz), essentially a neutral vowel. In many of the audio signals recorded from the singer, the fundamental frequency, *f*_0_, (vibratory frequency of the vocal folds) was typically about 150 Hz. He then appeared to enhance one of the harmonics in the approximate range of 8*f*_0_ … 12*f*_0_. Taking the 12th harmonic (12 × 150 = 1800 Hz) as an example target frequency (dashed line in the frequency response shown in Fig. 11a), the area function *A*_0_(*x*) was iteratively perturbed by the acoustic-sensitivity algorithm described in (***Story, 2006)*** until *F*_2_ and *F*_3_ converged on 1800 Hz and became a single formant peak in the frequency response. Additional details on the perturbation process leading into Fig.5 are detailed in the *Appendix*.

## Acknowledgments

A heartfelt thank you to Huun Huur Tu, without whom this study would not have been possible. Input/suggestions from Ralf Schlueter, Dorothea Kolossa, Chris Rozell, & Tuomas Virtanen are gratefully acknowledged. Support from York University, the Fields Institute for Research in Mathematical Sciences, and the Kavli Institute of Theoretical Physics are also gratefully acknowledged. C.B. was supported by Natural Sciences and Engineering Research Council of Canada (NSERC) Grant RGPIN-430761-2013.

## Appendix 1

The appendix contains supporting information for the document *Overtone focusing in Tuvan throat singing* by Bergevin et al.^*a*^. To provide a benchmark, we provide an overview of the several different sections present. First (**Methodological considerations**), we include several methodological components associated with the quantitative analysis of the waveforms, helping illustrate different approaches towards characterizing the acoustic data and rationale underlying *control* measures. Second (**Additional waveform analyses**), we include additional plots to support results and discussion in the main text. For example, different spectrograms are presented, as are analyses for additional waveforms. This section also helps provide additional context for a second independent focused state. The third section (**Additional modeling analysis figures**) details theoretical components leading into the results of the computational model and how the MRI data constrain the key parameters, so to help justify arguments surrounding the notion of formant merging. Fourth (**Instability in focused state**), some speculative discussion and basic modeling aspects are presented with regard to the notion of instabilities present in motor control of the focused state. In the fifth section (**Cochlear tonotopy**), basic aspects underlying cochlear tonotopy are presented and how such relates back to the observation that the salient perceptual focused state sits in the region of 1–2 kHz. Sixth (**Additional MRI analysis figures**), images stemming from the MRI data are presented. Lastly, the final three sections detail accessing the acoustic waveforms, MRI data files, and waveform analysis (Matlab-based) software via an online repository.

### Methodological considerations

#### Overtone & formant tracking

To facilitate quantification of the waveforms, we custom-coded a peak-picking/tracking algorithm to analyze the time-frequency representations produced by the spectrogram. Figure 1 shows an example of the tracking of the overtones (red dots) and formants (grayscale dots; intensity coded by relative magnitude as indicated by the colorbar). This representation provides an alternative view (compared to Fig.1) to help demonstrate that, by and large, the overtone structure is highly consistent throughout while the formant structure varies significantly across the transition.

**Appendix 1 Figure 1.**
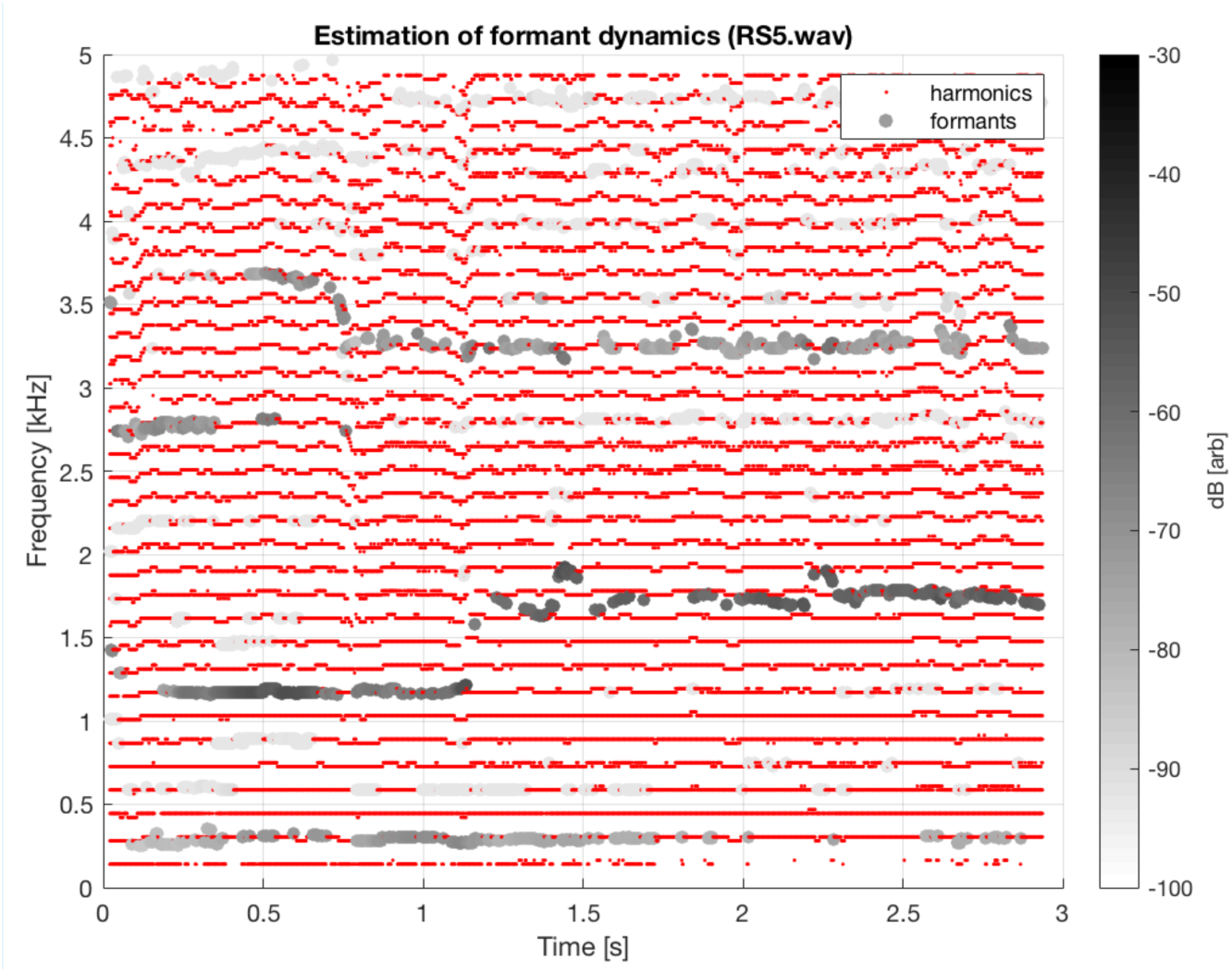
Same as Fig.1 (middle left panel; subject T2, same sound file as shown in middle panel of Fig.1), except with overtones and estimated formant structure tracked across time.

#### Quantifying Focused States

Figures 2 and 3 show calculation of the energy ratio *e*_*R*_ used as a means to quantify the degree of focus. For Fig.2, the waveforms are the same as those shown in Fig.1 (those with slightly different axis limits). In general, we found that *e*_*R*_(1, 2) provided a clear means to distinguish the focused state, as values were close to zero prior to the transition and larger/sustained beyond. Singer T2 was somewhat of an exception. Figure 3 is for singer T2, using the same file (i.e., the transition point into the focused state there between 6 and 7 s is the same as that shown in the middle panel of Fig.1), but with an expanded timescale to illustrate the larger *e*_*R*_ values prior to the transition. This is due to the relatively large amount of energy present between 2.5–4 kHz.

We also explored *e*_*R*_(1, 2) values in a wide range of phonetic signals, such as child & adult vocalizations, other singing styles (e.g., opera), non-Tuvan singers (e.g., Westerners) performing “overtone singing”, and older recordings of Tuvan singers. In general, it was observed that *e*_*R*_(1, 2) was relatively large and sustained across time for focused overtone song, whereas the value was close to zero and/or highly transient for other vocalizations.

**Appendix 1 Figure 2.**
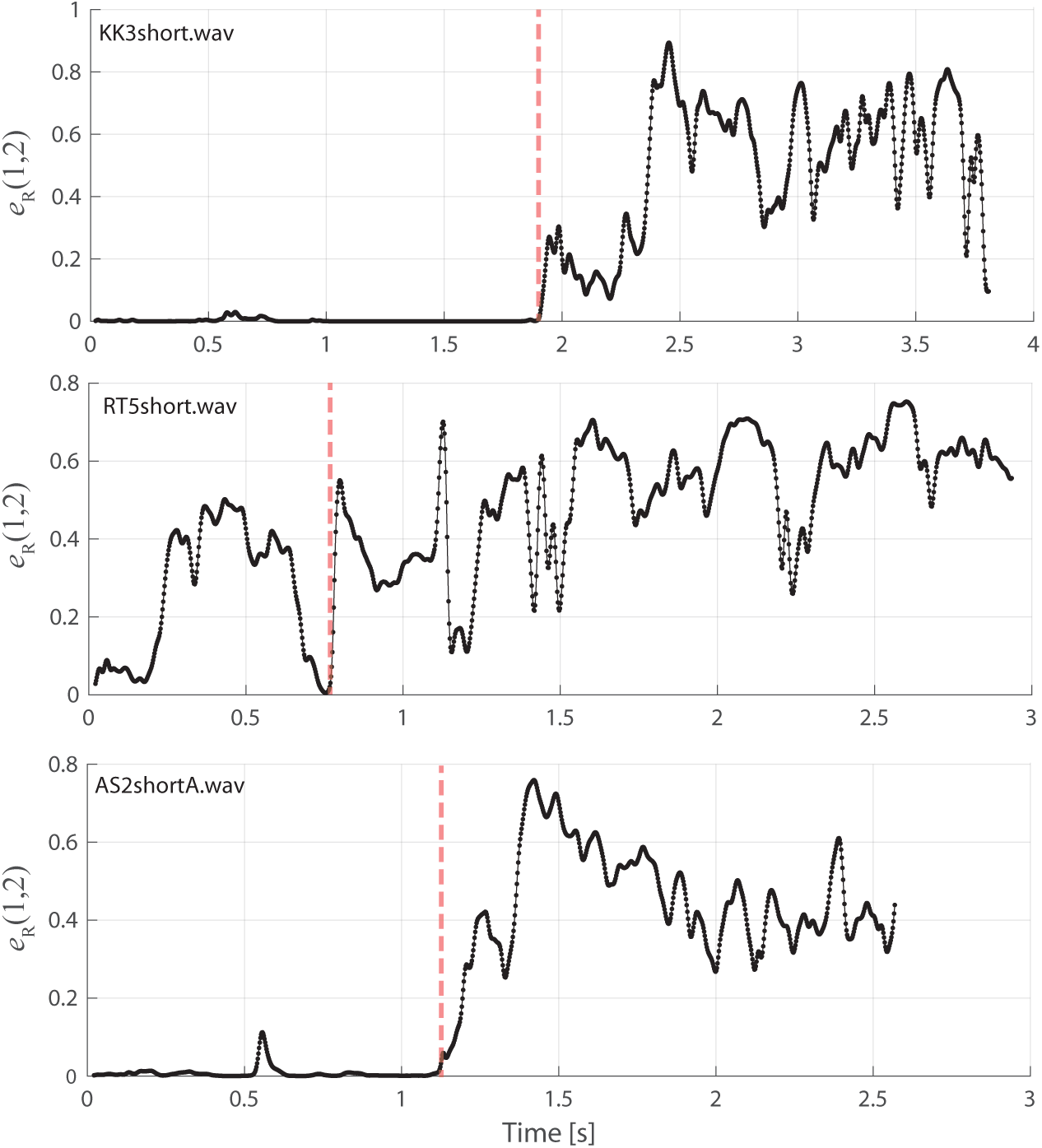
Same data/layout as in Fig.1 but now showing *e*_*R*_(1, 2) as defined in the Methods. That is, these plots show the energy ratio focused between 1–2 kHz. Vertical red dashed lines indicate approximate time of transition into the focused state. An expanded timescale is also shown for singer T2 (middle panel) is also shown in Fig.3.

**Appendix 1 Figure 3.**
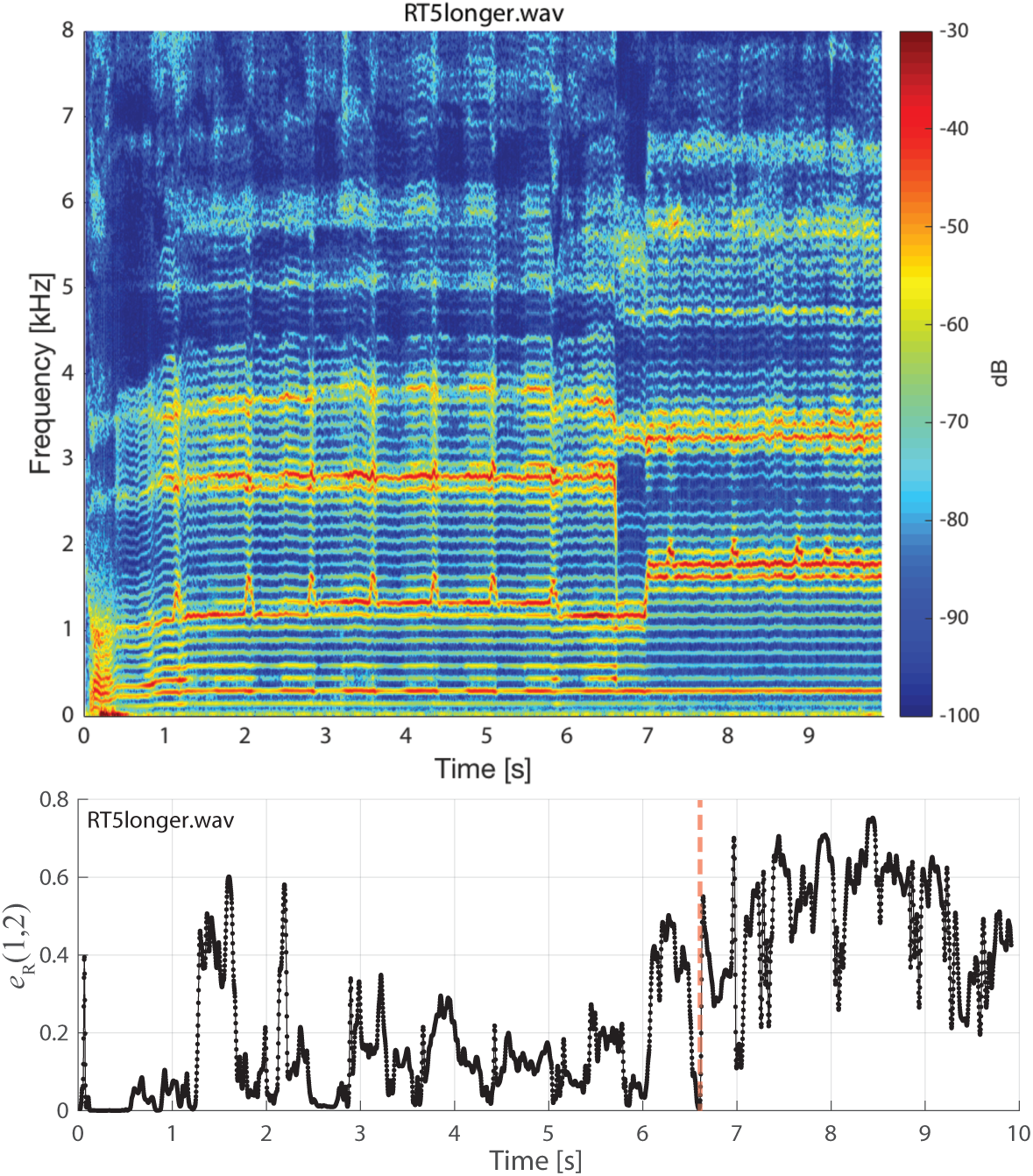
Similar to Fig.2 for singer T2 (middle panel), except an expanded time scale is shown to demonstrate the earlier dynamics as this singer approaches the focused state (see T2_5longer.wav).

#### Control measurements

The waveforms from the Tuvan singers provide an intrinsic degree of *control* (i.e., voicing not in the focused state). Similar in spirit to Fig.1, Fig.4 shows the spectra prior to the transition into the focused state. While relatively narrow harmonics can be observed, they tend to occur below 1 kHz. Such is consistent with our calculations of *e*_*R*_(1, 2): prior to a transition into a focused state, this value is close to zero. The exception is singer T2, who instead shows a relatively large amount of energy about 1.8-3 kHz that may have some sort of masking effect (see Discussion in main text, as well as Sec.**??**). Additionally, Tuvan singer T4 using a non-Sygyt (Fig.5) can also effectively be considered a “control”.

**Appendix 1 Figure 4.**
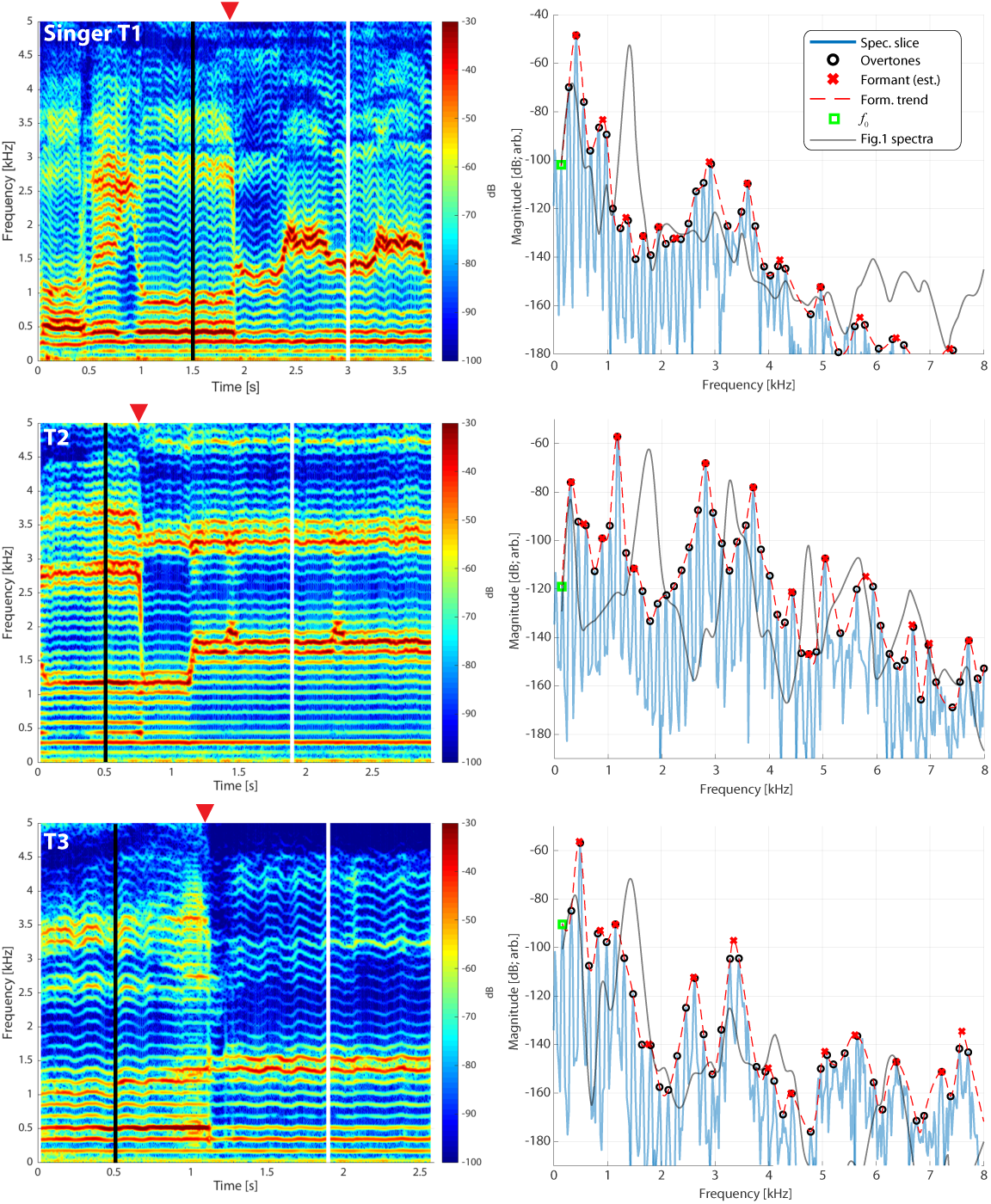
Stemming directly from Fig.1, the right-hand column now shows a spectrum from a time point prior to transition into the focused state (as denoted by the vertical black lines in the left column). The shape of the spectra from Fig.1 is also included for reference.

### Additional waveform analyses

#### Other spectrograms

Figure 5 shows spectrogram from singer T4 (T4_shortA.wav) singing in non-Sygyt style. While producing a distinctive sound, note the relative lack of energy above approximately 1 kHz. Figure 6 shows spectrogram from singer T2 (T2_5.wav) over a longer timescale than that shown in Fig.1. Similarly for Fig.7, but for singer T1. Both these plots provide a spectral-temporal view of how the singer maintain and modulate the song over the course of a single exhalation. Note both the sudden transitions into different maintained pitches, as well as the briefer transient excursions.

**Appendix 1 Figure 5.**
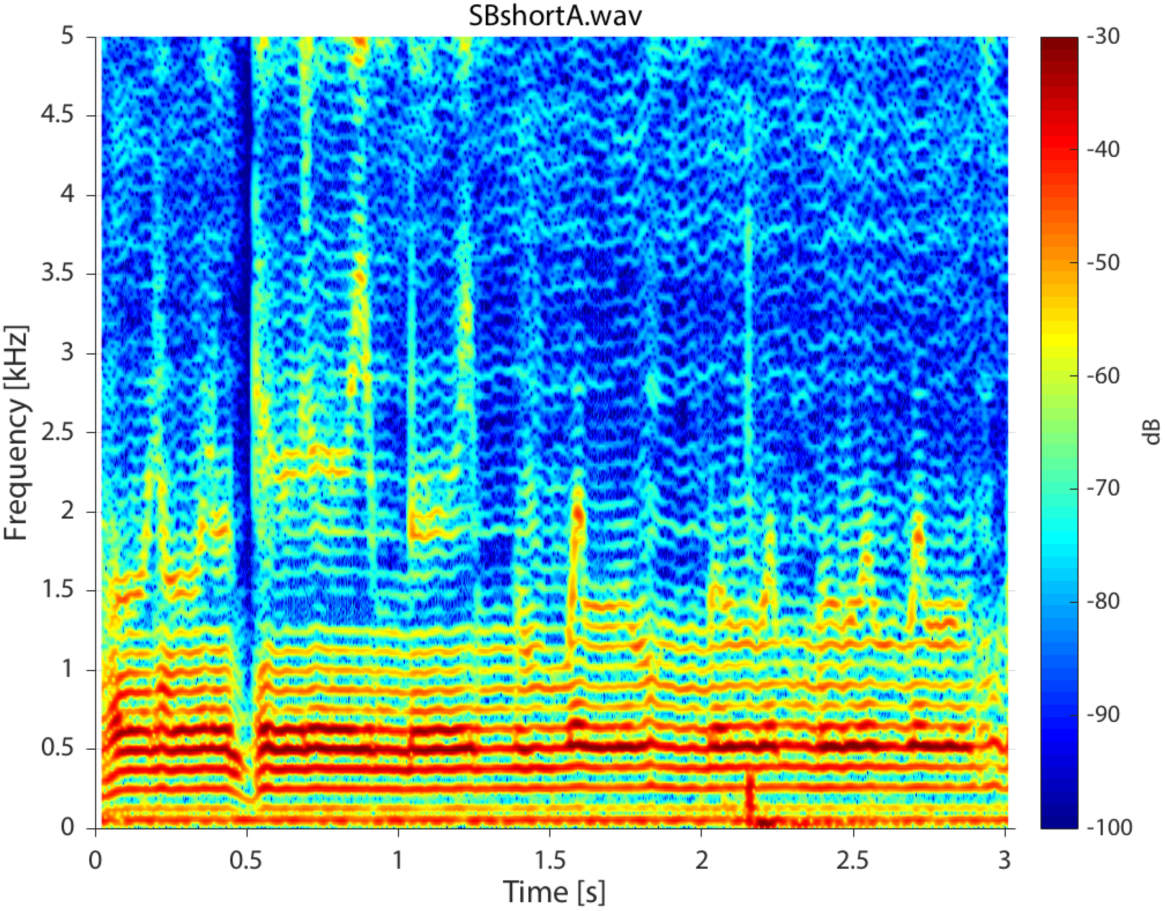
Spectrogram for singer T4 singing in non-Sygyt style (first song segment of T2_4shortA.wav sound file). For the spectrogram, 4096 point windows were used for the FFT with 95% fractional overlap and a Hamming window.

**Appendix 1 Figure 6.**
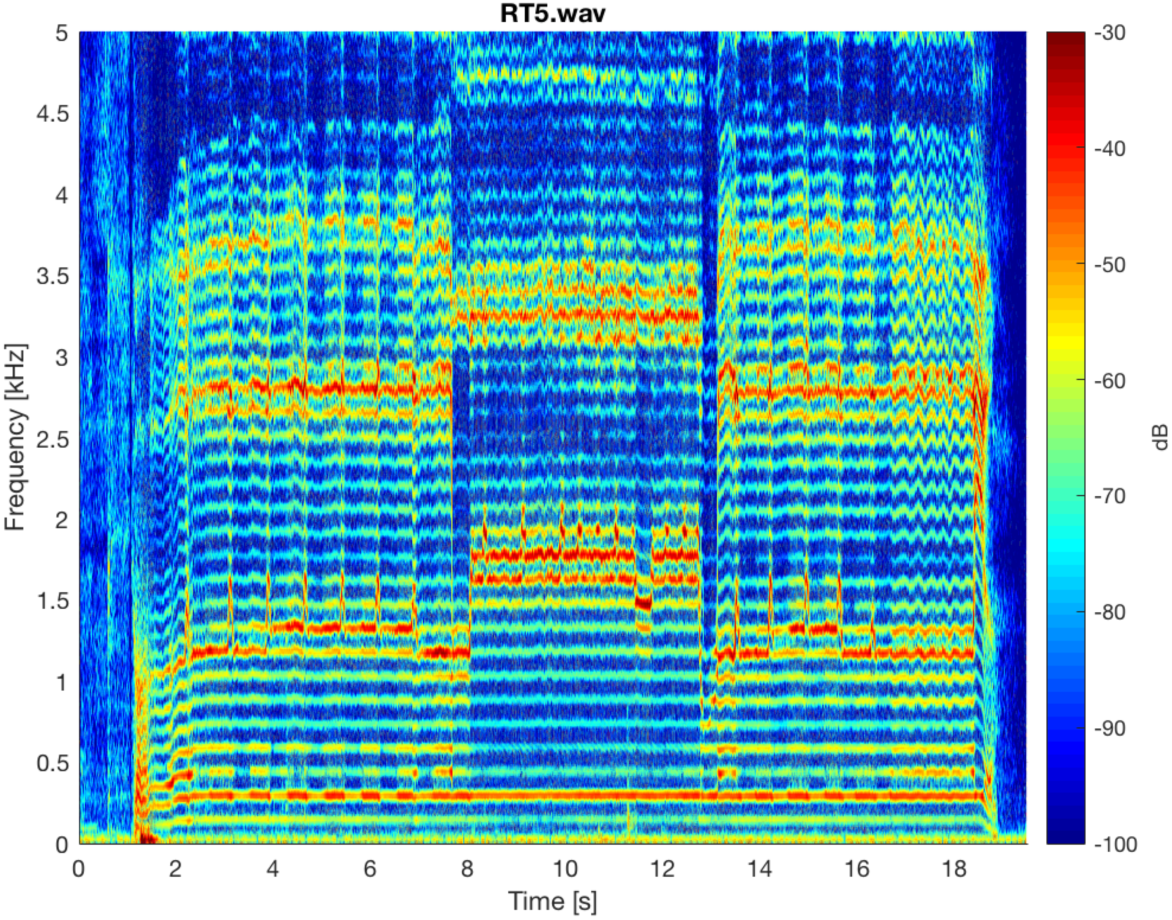
Spectrogram of entire T2_5.wav sound file. Sample rate was 96 kHz. The same analysis parameters were used as in Fig.5.

**Appendix 1 Figure 7.**
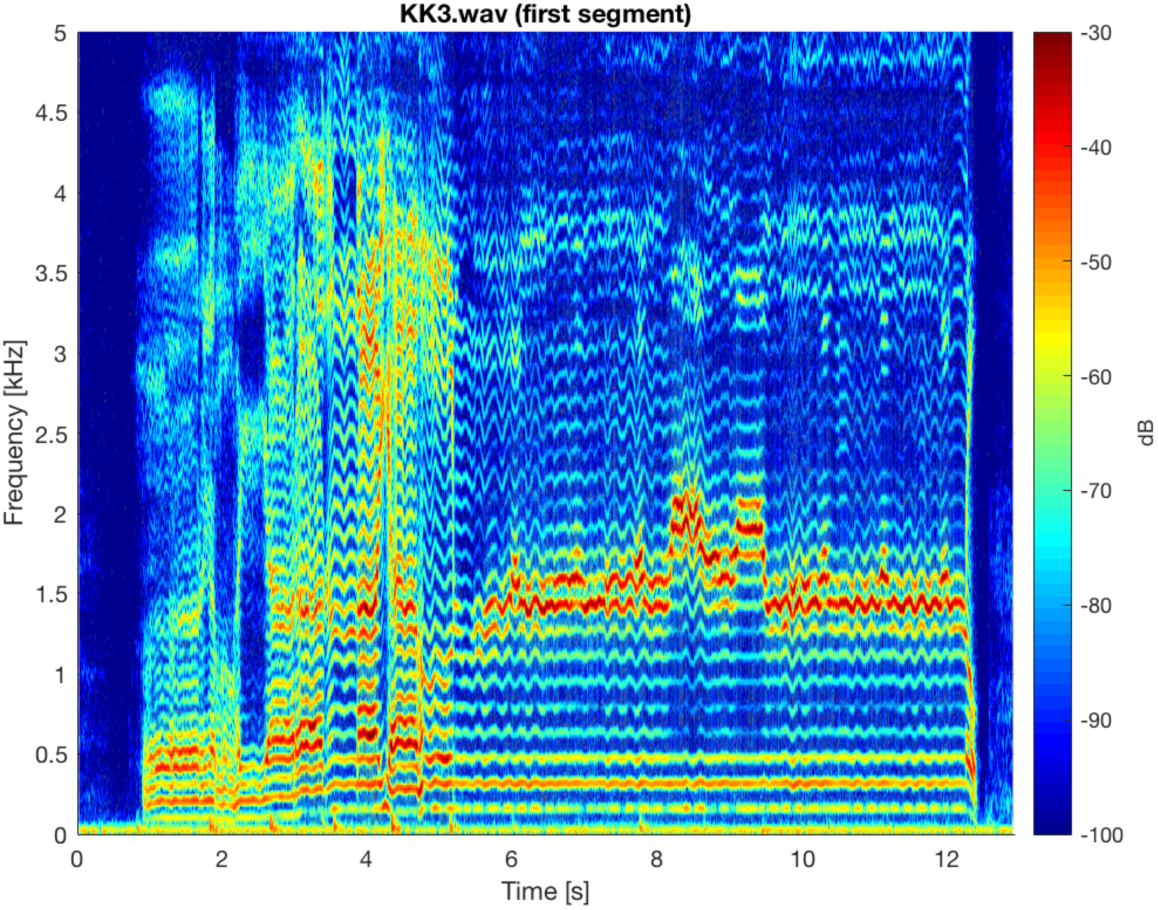
Spectrogram of first song segment of T1_3.wav sound file. The same analysis parameters were used as in Fig.5.

#### Second independent focused state

Figure 8 shows another example of a transition in Sygyt-style song for singer T2, clearly showing a second focused state about 3–3.5 kHz. Two aspects merit highlighting. First, the spectral peaks are not harmonically-related: At *t* = 4.5 s, the first focused state is at kHz and the other at 3.17 kHz (far from an expected to 2.72 kHz). Second, during the singer-induced pitch change at 3.85 s, the two peaks do not move in unison. While not ruling out correlations between the two focused states, these observations suggest that they are not simply nor strongly related to one another.

**Appendix 1 Figure 8.**
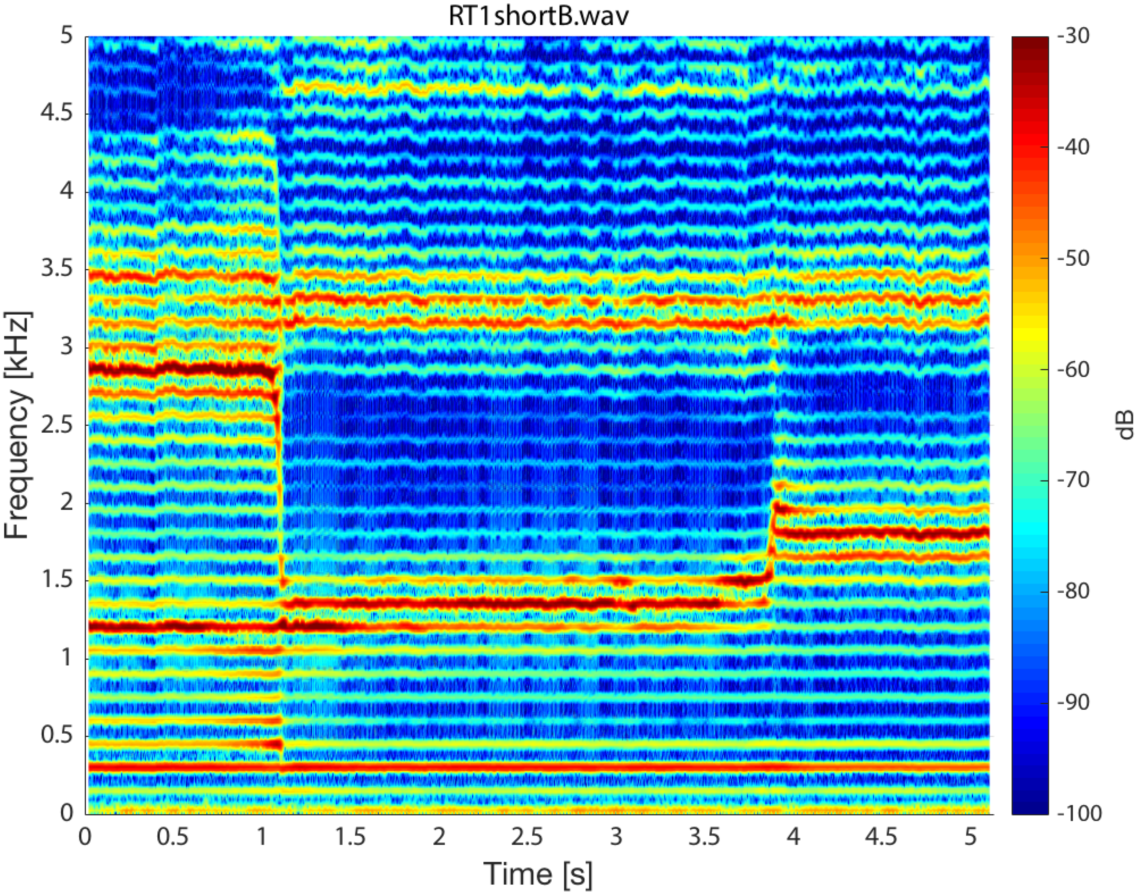
Singer T2 about transition into a focused state. Note that while the first focused state transitions from approximately 1.36 to 1.78 kHz, the second state remains nearly constant, decreasing only slightly from 3.32 to 3.17 kHz. (T2_1shortB.wav)

#### Pressed transition

Figure 9 shows a spectrogram and several spectral slices for the sound file where the voicing was “pressed” (***Adachi and Yamada, 1999***; ***Edmondson and Esling, 2006)*** prior to the transition into the focused state. That is, prior to the 1.8 s mark, voicing is relatively normal. But after that point (prior to the transition into the focused state around 5.4 s, substantial energy appears between 2–4 kHz along with a degree of vibratto. Note however that there is no change to the overall overtone structure (e.g., no emergence of subharmonics). The spectrum at *t* = 4.0s, prior to the transition, provides a useful comparison back to (***Levin and Süzükei, 2006)***. Specifically, one particular overtone is singled out and highly focused, yet the broadband cluster of overtones about 2.5–4 kHz effectively mask it. It is not until about the 5.4 s mark when those higher overtones are also brought into focus that a salient perception of the Sygyt-style emerges.

**Appendix 1 Figure 9.**
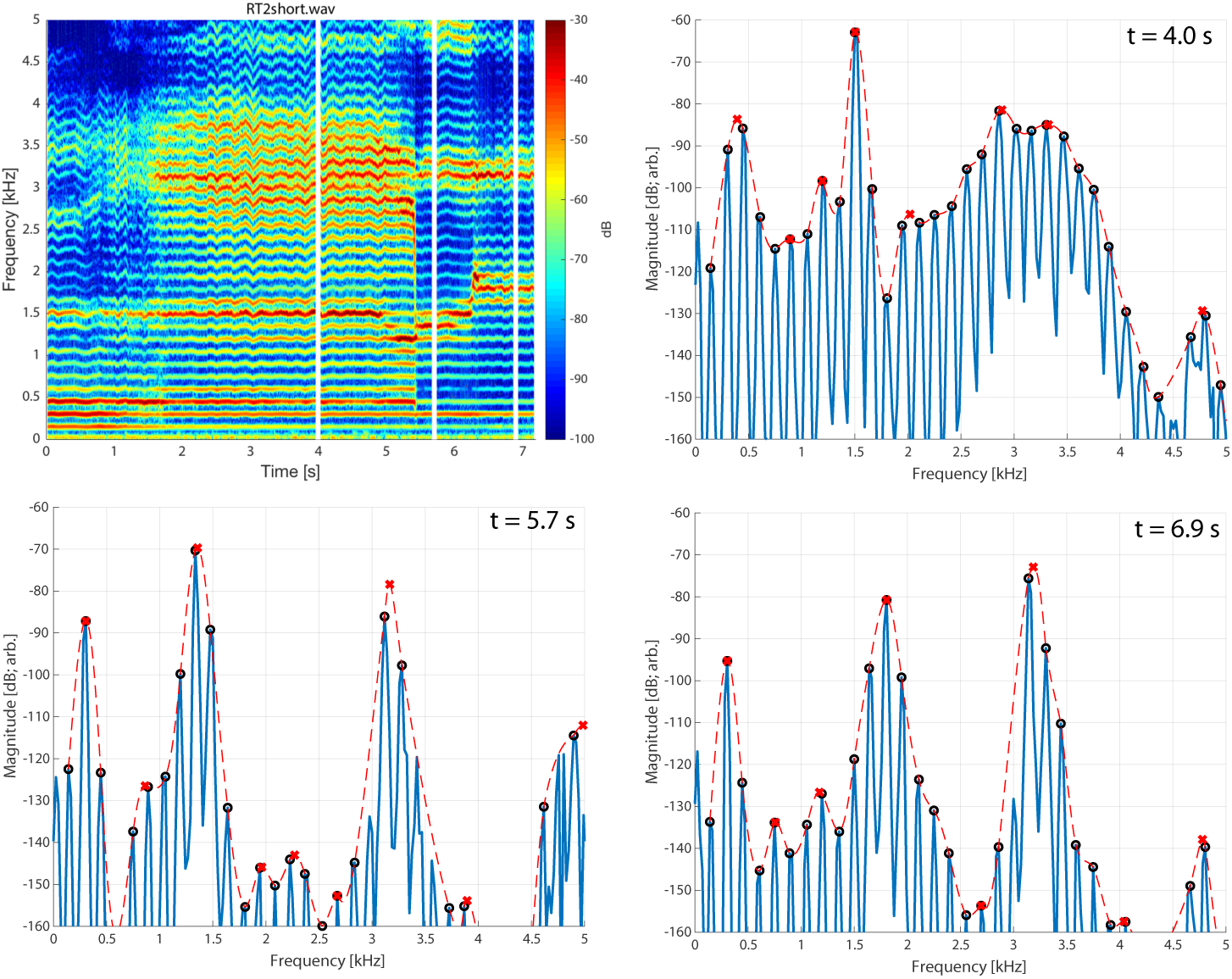
Spectrogram of singer T2 exhibiting pressed voicing heading into transition to focused state (T2_2short.wav).

### Additional modeling analysis figures

The measurement of the cross-distance function (as described in the *Methods*), along with calculation of the frequency response from an estimate of the area function, suggested that constrictions of the vocal tract in the region of the uvula and alveolar ridge may play a significant role in controlling the spectral focus generated by the convergence of *F*_2_ and *F*_3_. Assuming that an overtone singer configures the vocal tract to deliberately merge these two formants such that, together, they enhance the amplitude of a selected harmonic of the voice source, the aim was to investigate how the vocal tract can be systematically shaped with precisely-placed constrictions and expansions to both merge *F*_2_ and *F*_3_ into a focused cluster *and* move the cluster along the frequency axis to allow for selection of a range of voice harmonics.

Shown in Fig. 11b is the same area function as Fig. 11a (see Methods) but plotted by extending the equivalent radius of each cross-sectional area, outward and inward, along a line perpendicular to the centerline measured from the singer (see Fig. 4a), resulting in an inner and outer outline of the vocal tract shape as indicated by the thick black lines. The measured centerline is also shown in the figure, along with anatomic landmarks. Since this does not represent a true midsagittal plane it will be referred to here as a *pseudo-midsagittal* plot (***Story et al., 2001)***.

Fig. 12a shows the new area function and frequency response generated by the perturbation process, whereas the pseudo-midsagittal plot is shown in Fig. 12b. Relative to the shape of *A*_0_(*x*) (shown as the thin gray line), the primary modification is a severe constriction imposed between 12.5-13.5 cm from the glottis, essentially at the alveolar ridge. Although the line thickness might suggest that the vocal tract is occluded in this region, the minimum cross-sectional area is 0.09 cm^2^. There is also a more moderate constriction at about 5 cm from the glottis, and a slight expansion between 7-10.5 cm from the glottis. The frequency response in upper panel of Fig. 12a demonstrates that the new area function was successful in driving *F*_2_ and *F*_3_ together to form a single formant peak centered at 1800 Hz that is at least 15 dB higher in amplitude than any of the other formants.

Exactly the same process was used to generate area functions for which *F*_2_ and *F*_3_ converge on the target harmonic frequencies: 8*f*_0_, 9*f*_0_, 10*f*_0_, 11*f*_0_ = 1200, 1350, 1500, 1650 Hz. The results, along with those from the previous figure for 12*f*_0_, are shown in Fig. 13. The collection of frequency responses in the upper panel of Fig. 13b shows that *F*_2_ and *F*_3_ successfully converged to become one formant peak in each of the cases, and their locations on the frequency axis are centered around the specified target frequencies. The corresponding area functions in the lower panel suggests that the constriction between 12.5-13.5 cm from the glottis (alveolar ridge region) is present in roughly the same form for all five cases. In contrast, an increasingly severe constriction must be imposed in the region between 6-8.5 cm from the glottis (uvular region) in order to shift the target frequency (i.e., the frequency at which *F*_2_ and *F*_3_ converge) downward through progression of specified harmonics. Coincident with this constriction is a progressively larger expansion between 14-15.5 cm from the glottis which likely assists in positioning the focal regions of *F*_2_ and *F*_3_ downward. It can also be noted that the area function that generates a focus at 8*f*_0_ (1200 Hz; thinnest line) is most similar the one generated from the cross-distance measurements (i.e., Fig. 4c). In both, there are constrictions located at about 7.5 cm and 13 cm from the glottis; the expansions in the lower pharynx and oral cavity are also quite similar. The main difference is the greater expansion of the region between 8-13 cm from the glottis in the acoustically-derived area function.

Based on the results, a mechanism for controlling the enhancement of voice harmonics can be proposed: The degree of constriction near the alveolar ridge in the oral cavity, labeled *C*_0_ in Fig. 13, controls the proximity of *F*_2_ and *F*_3_ to each other, whereas the degree of constriction near the uvula in the upper pharynx, *C*_*p*_, controls the frequency at which *F*_2_ and *F*_3_ converge (the expansion anterior to *C*_0_ may also contribute). Thus, an overtone singer could potentially “play” (i.e., select) various harmonics of the voice source by first generating a tight constriction in the oral cavity near the alveolar ridge to generate the focus of *F*_2_ and *F*_3_, and then modulating the degree of constriction in the uvular region of the upper pharynx to position the focus on a selected harmonic.

This proposed mechanism of controlling the spectral focus is supported by observation of vocal tract changes based on dynamic MRI data sets. Using this approach midsagittal movies of the Tuvan singer were acquired where each image represented approximately 275 ms. Shown in Fig. **??** is a comparison of vocal tract configurations derived with the acoustic-sensitivity algorithm (middle panels) to image frames from an MRI-based movie (upper panels) associated with the points in time indicated by the vertical lines superimposed across the waveform and spectrogram in the lower part of the figure. The image frames were chosen such that they appeared to be representative of the singer placing the spectral focus at 8*f*_0_ (left) and 12*f*_0_ (right), respectively, based on the evidence available in the spectrogram. The model-based vocal tract shape in the upper left panel, derived for a spectral focus of 8*f*_0_ (1200 Hz), exhibits a fairly severe constriction in the uvular region, similar to the constrictive effect that can be seen in the corresponding image frame (middle left). Likewise, the vocal tract shape derived for a spectral focus of 12*f*_0_ (1800 Hz) (upper right) and the image frame just below it both demonstrate an absence of a uvular constriction. Thus, the model-based approach generated vocal tract shapes that appear to possess characteristics similar to those produced by the singer, and provides support for the proposed mechanism of spectral focus control.

**Appendix 1 Figure 10.**
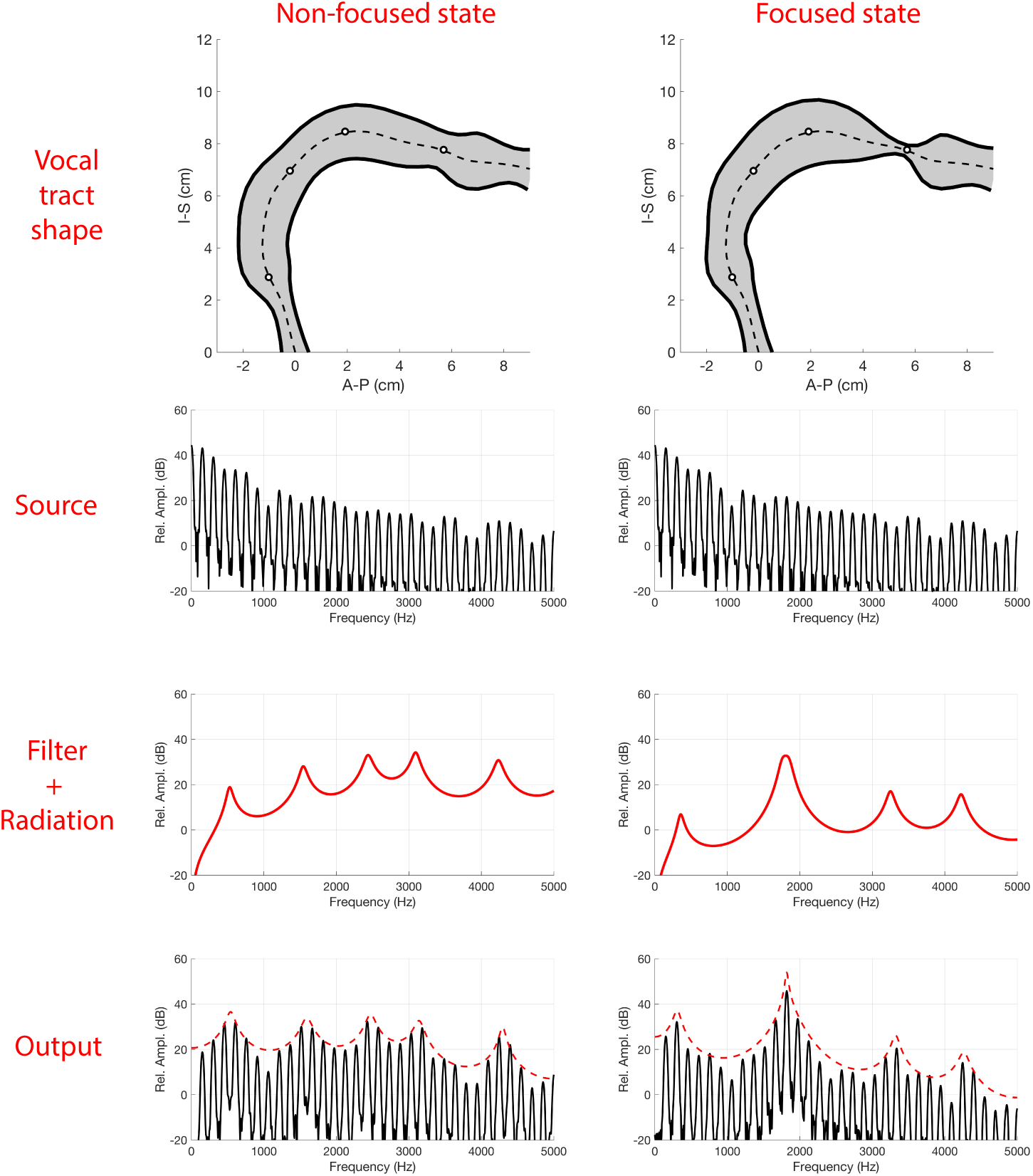
Overview of source/filter theory, as originally motivated by (**?**, stevens2000acoustic) Left column shows normal phonation, while the right indicates one example of a focused state.

**Appendix 1 Figure 11.**
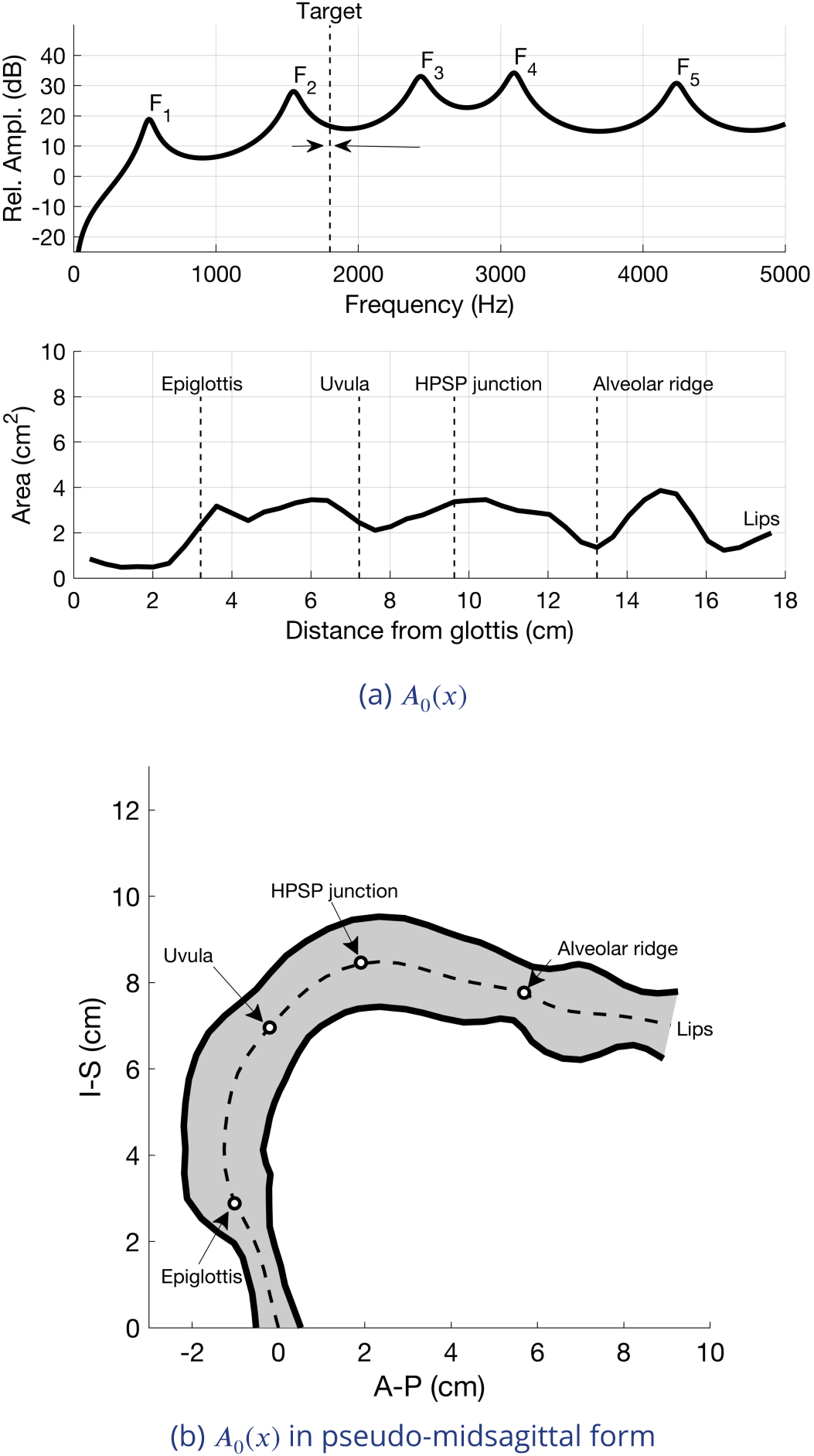
Setup of the baseline vocal tract configuration used in the modeling study. (a) The area function is in the lower panel and its frequency response is in the upper panel. (b) The area function from (a) is shown as a pseudo-midsagittal plot (see text).

**Appendix 1 Figure 12.**
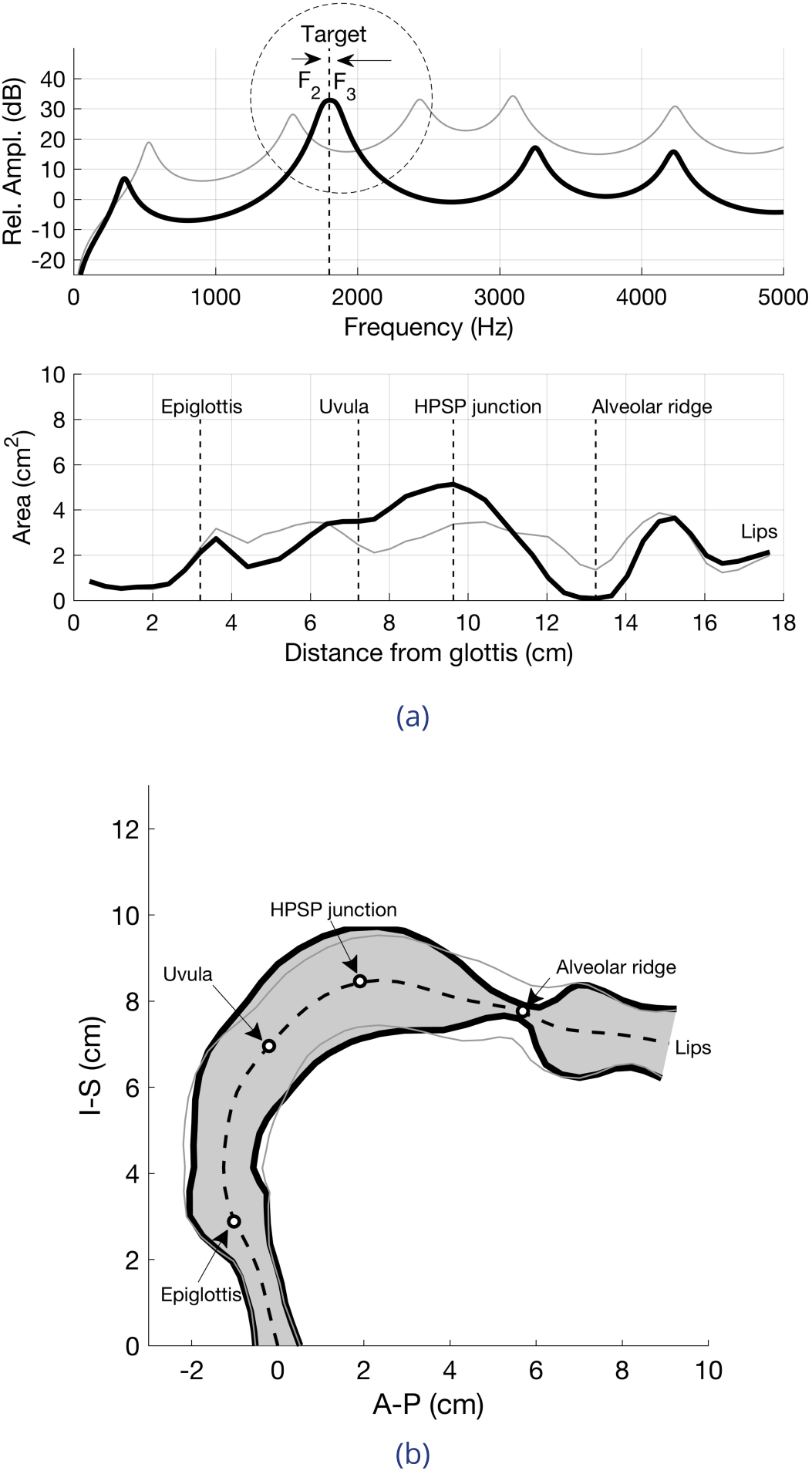
Results of perturbing the baseline area function *A*_0_(*x*) so that *F*_2_ and *F*_3_ converge on 1800 Hz. (a) perturbed area function (thick black) and corresponding frequency response; for comparison, the baseline area function is also shown (thin gray). The frequency response shows the convergence of *F*_2_ and *F*_3_ into one high amplitude peak centered around 1800 Hz. (b) Pseudo-misagittal plot of the perturbed area function (thick black) and the baseline area function (thin gray).

**Appendix 1 Figure 13.**
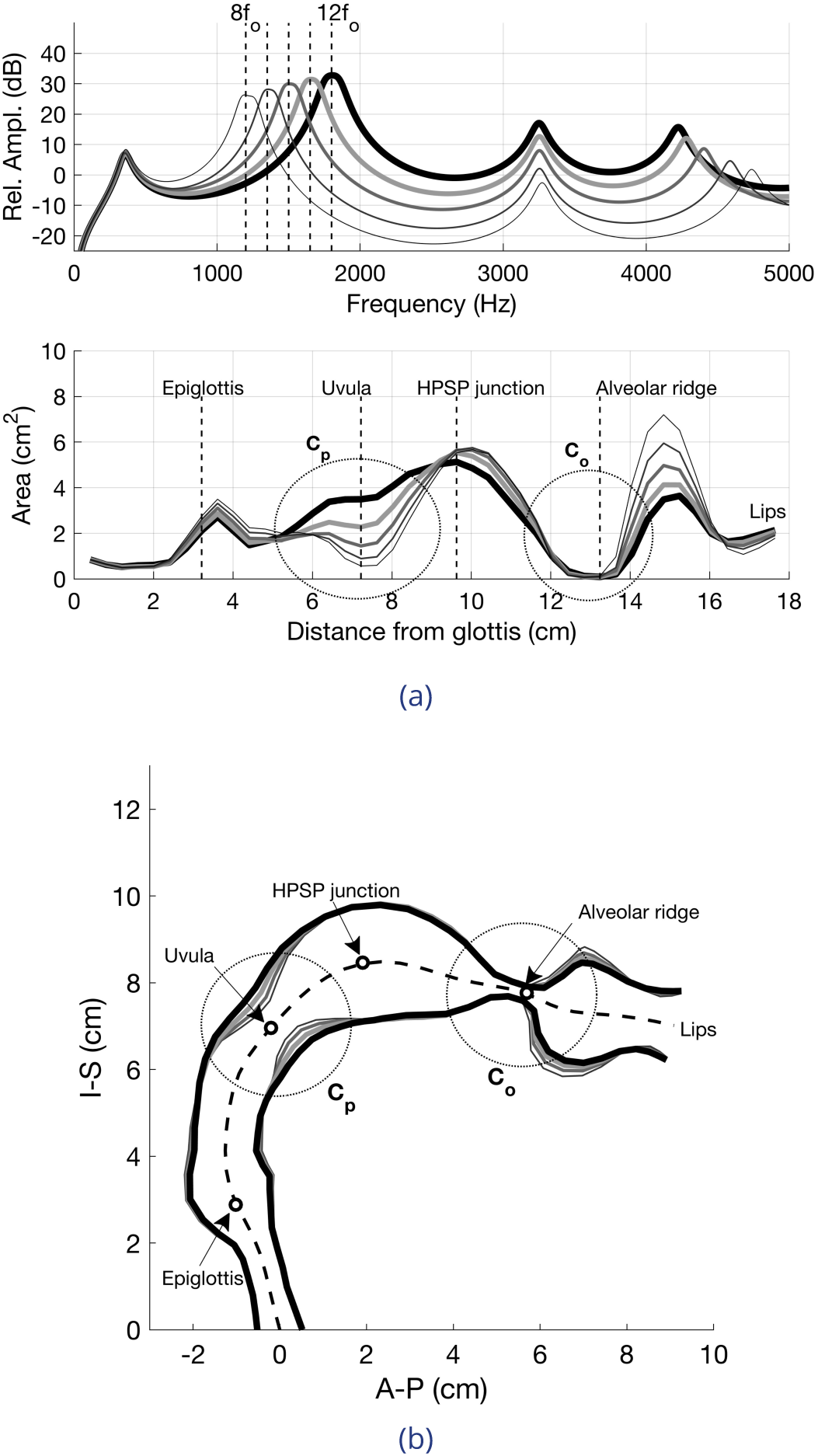
Results of perturbing the baseline area function *A*_0_(*x*) so that *F*_2_ and *F*_3_ converge on 1200, 1350, 1500, 1650, and 1800 Hz, respectively. (a) perturbed area functions and corresponding frequency responses; line thicknesses and gray scale are matched in the upper and lower panels. (b) Pseudo-misagittal plot of the perturbed area functions. The circled regions (dotted) denote constrictions that control the proximity of *F*_2_ and *F*_3_ to each other and the frequency at which they converge.

#### Second focused state

Given that singer T2 was the subject for the MRI scans and uniquely exhibited a second focused state [e.g., Fig.8], the model was also utilized to explore how multiple states could be achieved. Two possibilities appear to be the sharpening of formant F4 alone, or the merging of F4 and F5 (Fig.14). However, it is unclear how reasonable those vocal tract configurations may be and further study is required.

**Appendix 1 Figure 14.**
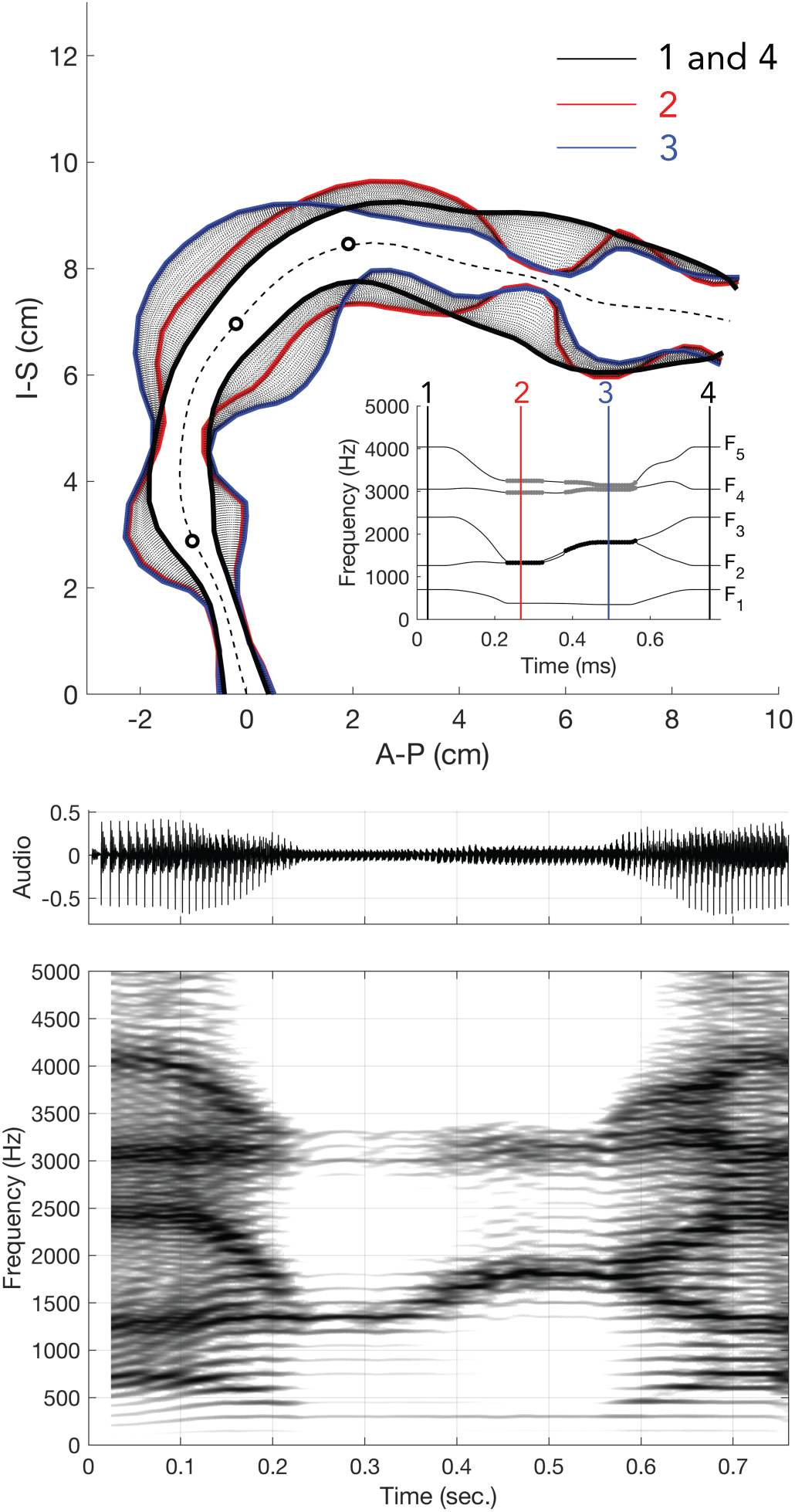
Similar to Fig.5, but additional manipulations were considered to create a second focused state by merging F4 and F5, as exhibited by singer T2 (see middle row in Fig.1). Additionally, the spectrogram shown here is from the model (not the singer’s audio). See also Fig.21 for connections back to dynamic MRI data.

#### Animations & synthesized song

These animations and audio clips demonstrate various quantitative aspects of the model.

- Animation (no sound) of vocal tract changes during transition into focused state and subsequent pitch changes – https://www.dropbox.com/s/hbtn44oaepyugyb/Medley_0to5_1cluster.mp4?dl=0
- Audio clip of simulated song – https://www.dropbox.com/s/4bgt0pj3pqk8nqs/Medley_0to5_1cluster_sim.wav?dl=0

### Instability in focused state

Figures 15 and 16 show that brief transient instabilities in the focused state can and do regularly occur. Specifically, it can be observed that there are brief transient lapses while the singer is maintaining the focused overtone condition, thereby providing insight into how focus is actively maintained. One possible explanation is control by virtue of biomechanical feedback, where the focused state can effectively be considered as an unstable equilibrium point, akin to balancing a ruler vertically on the palm of your hand. An alternative consideration might be that singers learn to create additional quasi-stable equilibrium points (e.g., Fig.17). The sudden transitions observed (Fig.1) could then be likened to two-person cheer-leading moves such as a “cupie”, where one person standing on the ground suddenly throws another up vertically and has them balancing atop their shoulders or upward-stretched hands. A simple proposed model for the transition into the focused state is shown in Fig.17. There, a stable configuration of the vocal tract would be the low point (pink ball). Learning to achieve a focused state would give rise to additional stable equilibiria (red ball), which may be more difficult to maintain. Considerations along these lines combined with a model for biomechanical control [e.g., ***Sanguineti et al. (1998)***] can lead to testable predictions specific to when a highly experienced singer is maintaining balance about the transition point into/out of a focused state (e.g., T2_4.wav audio file).

**Appendix 1 Figure 15.**
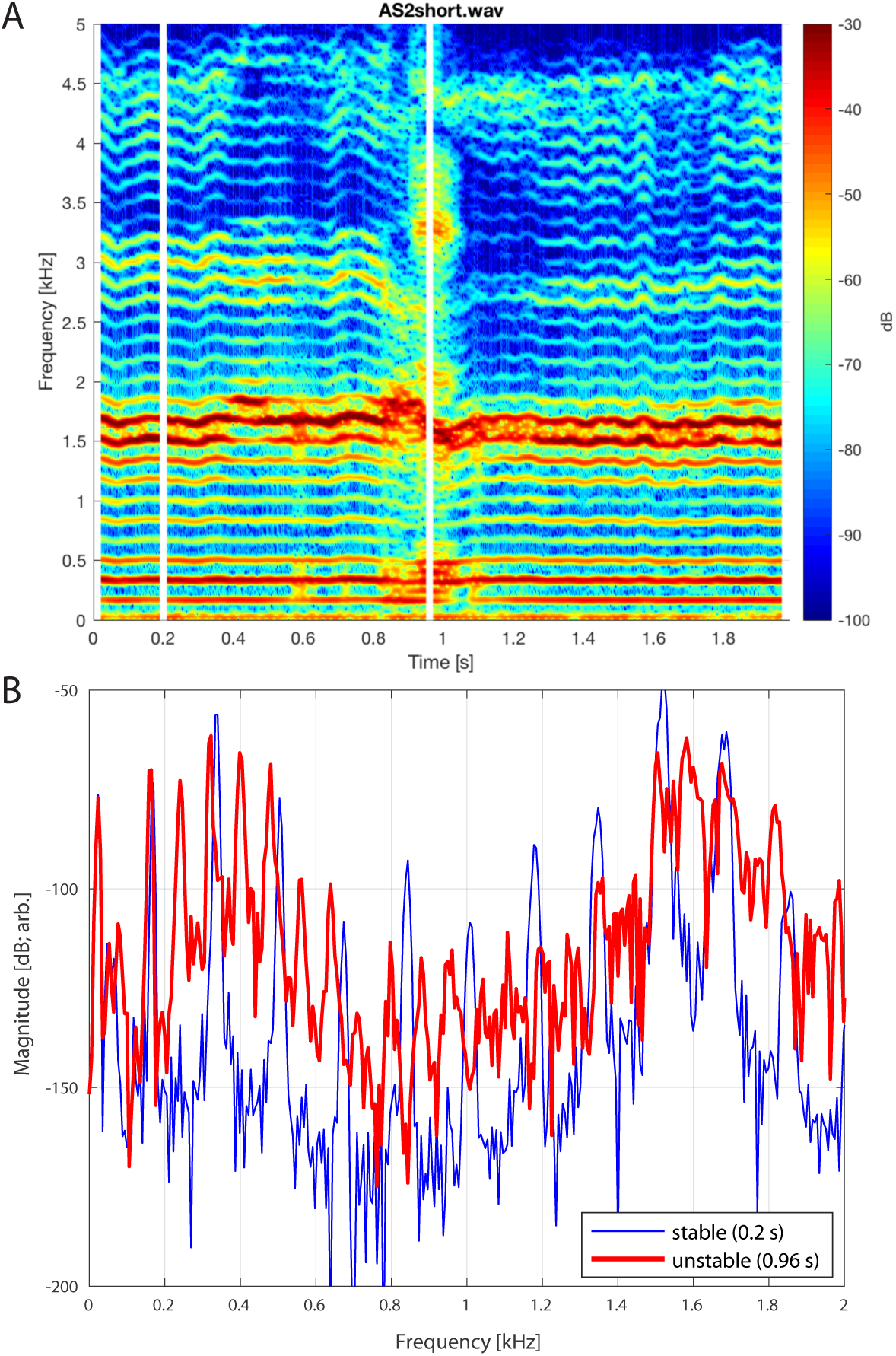
A – Spectrogram of singer T3 during period where the focused state briefly falters (T3_2shortB.wav, extracted from around 33 s mark of T3_2.wav). B – Spectral slices taken at two different time points (vertical white lines in A at 0.2 and 0.96 s), the latter falling in the transient unstable state. Note that while there is little change in *f*_0_ between the two periods (170 versus 164 Hz), the unstable period shows a period doubling such that the subharmonic (i.e., f0/2) and associated overtones are now present, indicative of nonlinear phonation.

**Appendix 1 Figure 16.**
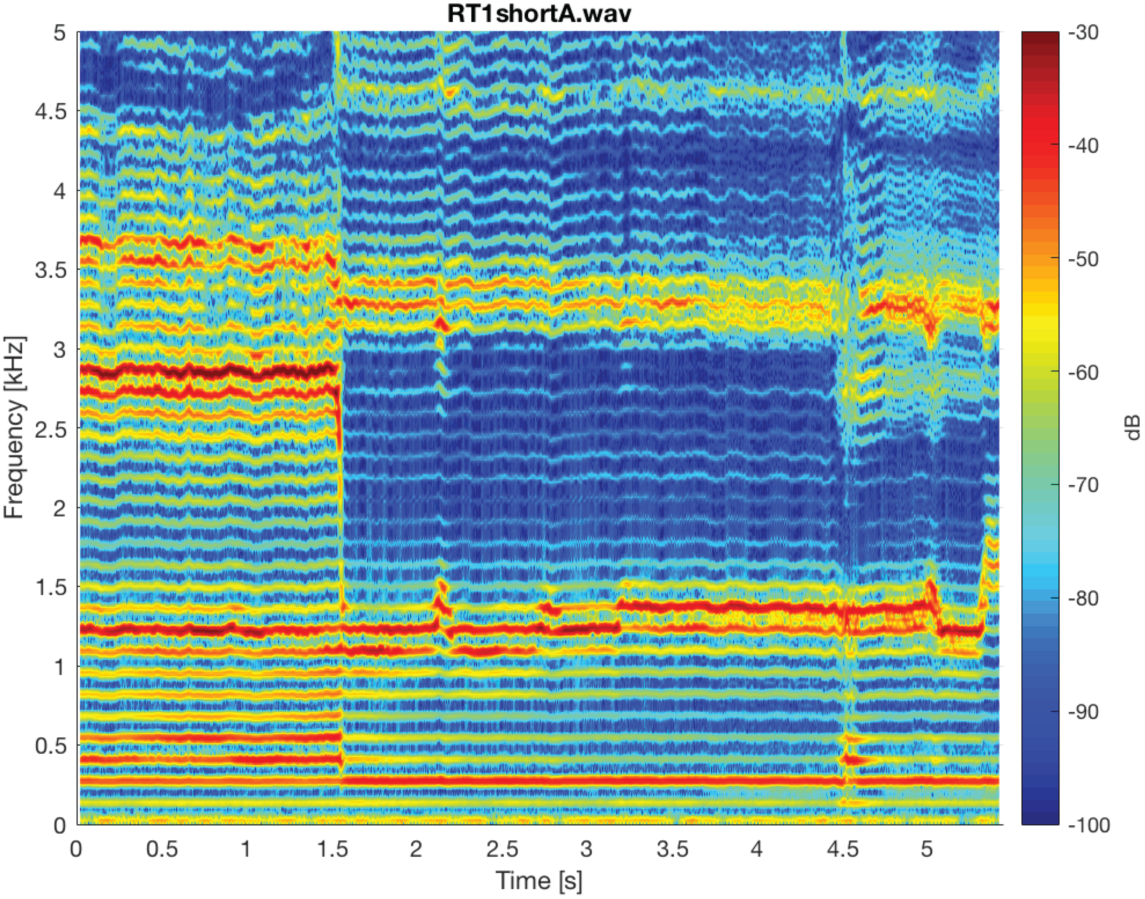
Spectrogram of singer T2 (T2_1shortA.wav) about a transition into a focused state. Note around 4.5 s, where there is a slight instability.

**Appendix 1 Figure 17.**
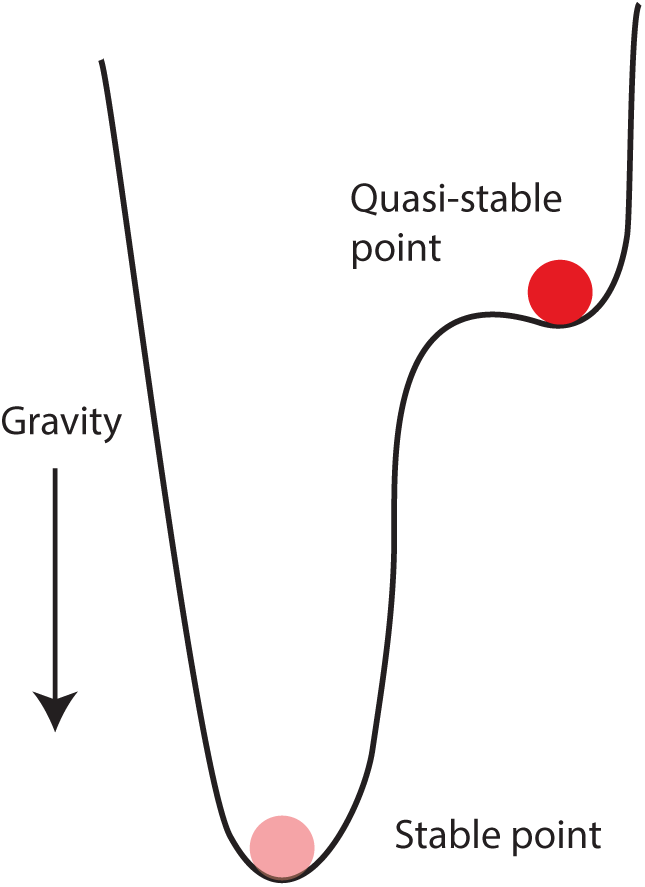
Schematic illustrating a simple possible mechanical analogy (ball confined to a potential well) for the transition into a focused state.

### Cochlear tonotopy

To help explain the rationale for how the Sygyt-style song may capitalize off a unique trait of the human cochlea, this section provides some basic aspects of human cochlear tonotopy [i.e., how characteristic frequency, CF, maps to spatial position along the length of the basilar membrane (BM) relative to the base, *x*]. While the precise form of the human tonotopic map is unknown, Fig.18 compares a simple exponential mapping with a commonly assumed form, the so-called *Greenwood map*. The simple exponential map is given by

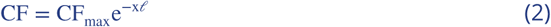

where 𝓁 is the “space constant” [mm] and CF_max_ [kHz] is the highest frequency represented on the BM. In Fig.18, we used CF_max_ = 20 and 𝓁 = 7. The Greenwood map (***Greenwood, 1990)*** is given by

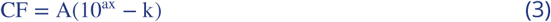

Commonly used parameters are *A* = 165.4, *a* = 0.06, and *k* = 0.88. Such allows the frequency range to span approximately 10 octaves from 20–20000 Hz. Note that the region of focused song (1–2 kHz, gray region in Fig.18) sits just basal to where the two types of map diverge, which corresponds also to the proposed 1 kHz “apical–basal transition CF” for humans (***Shera et al., 2010)***.

**Appendix 1 Figure 18.**
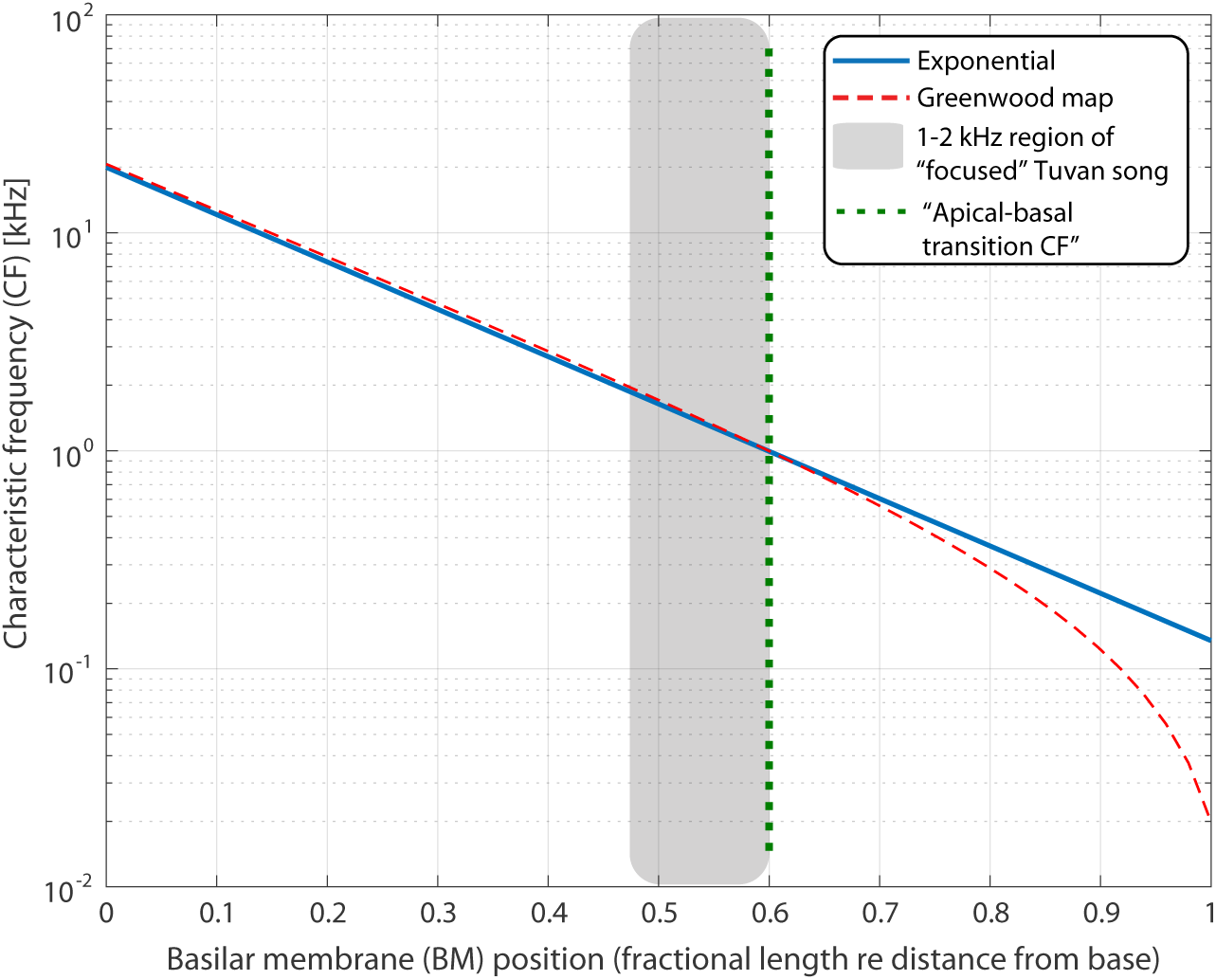
Estimates of the human tonotopic map. That is, the characteristic frequency (CF) as a function (see text) of position along the length of the cochlea relative to the base (*x*). Blue curve shows a simple exponential mapping, while the red curve indicates the “Greenwood map” (***Greenwood, 1990)***. Grey region indicates the 1–2 kHz octave range spanned by the focused Tuvan song and green dashed line the “apical–basal transition CF” (***Shera et al., 2010)***.

### Additional MRI analysis figures

#### Volumetric data

An example of the volumetric data (arranged as tiled midsagittal slices) is shown in Fig.19. Note that the NMR artifact due to the presence of a dental post is apparent lateralized to one side.

**Appendix 1 Figure 19.**
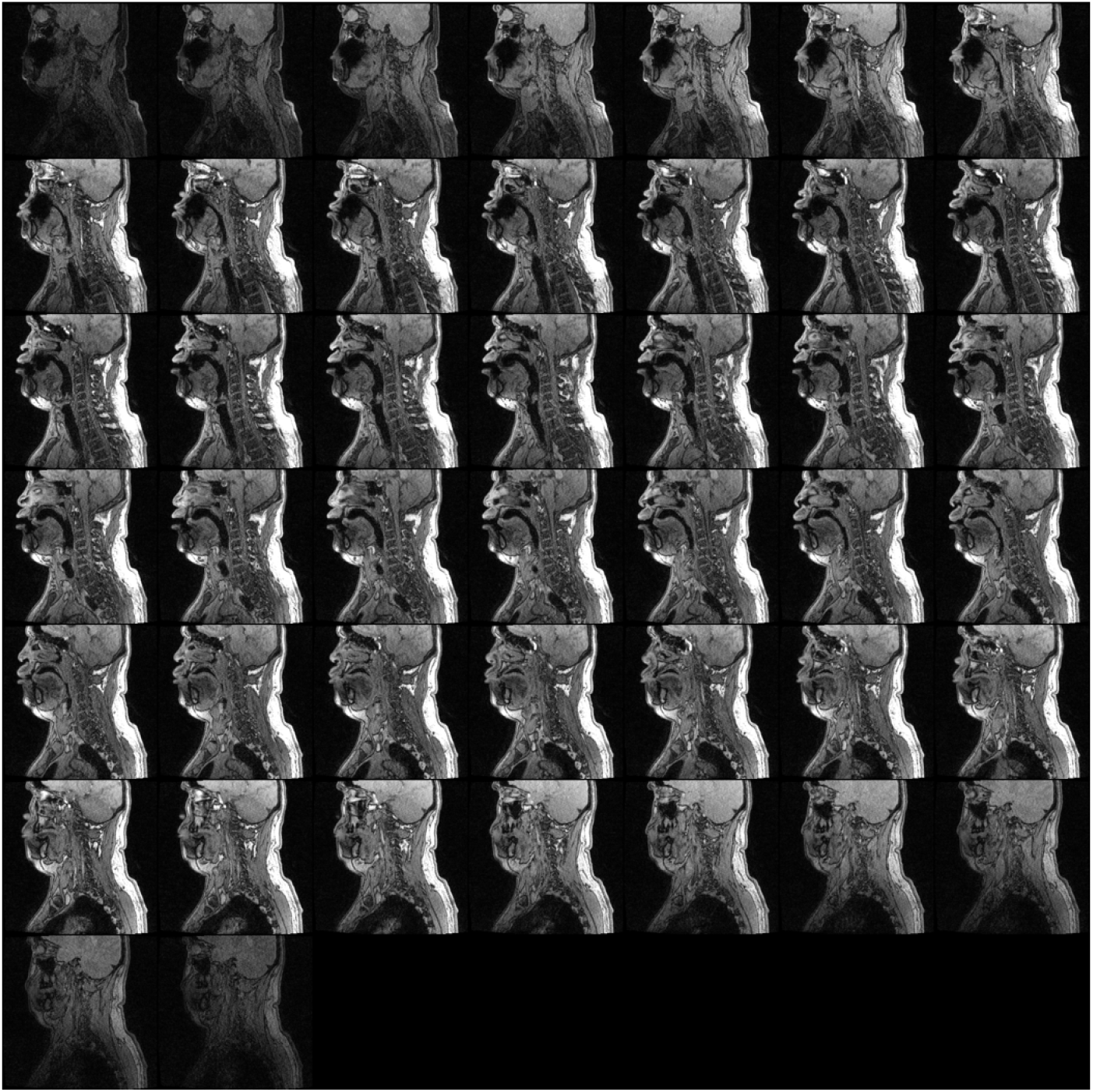
Mosaic of single slices from the volumetric MRI scan (Run3) of subject T2 during focused overtone state. Spectrogram of corresponding audio shown in Fig.20.

Figure 20 shows a spectrogram of audio segment (extracted from Run3Vsound.wav) associated with volumetric scan shown in Fig.19. Segments both with and without the scanner noise are shown.

**Appendix 1 Figure 20.**
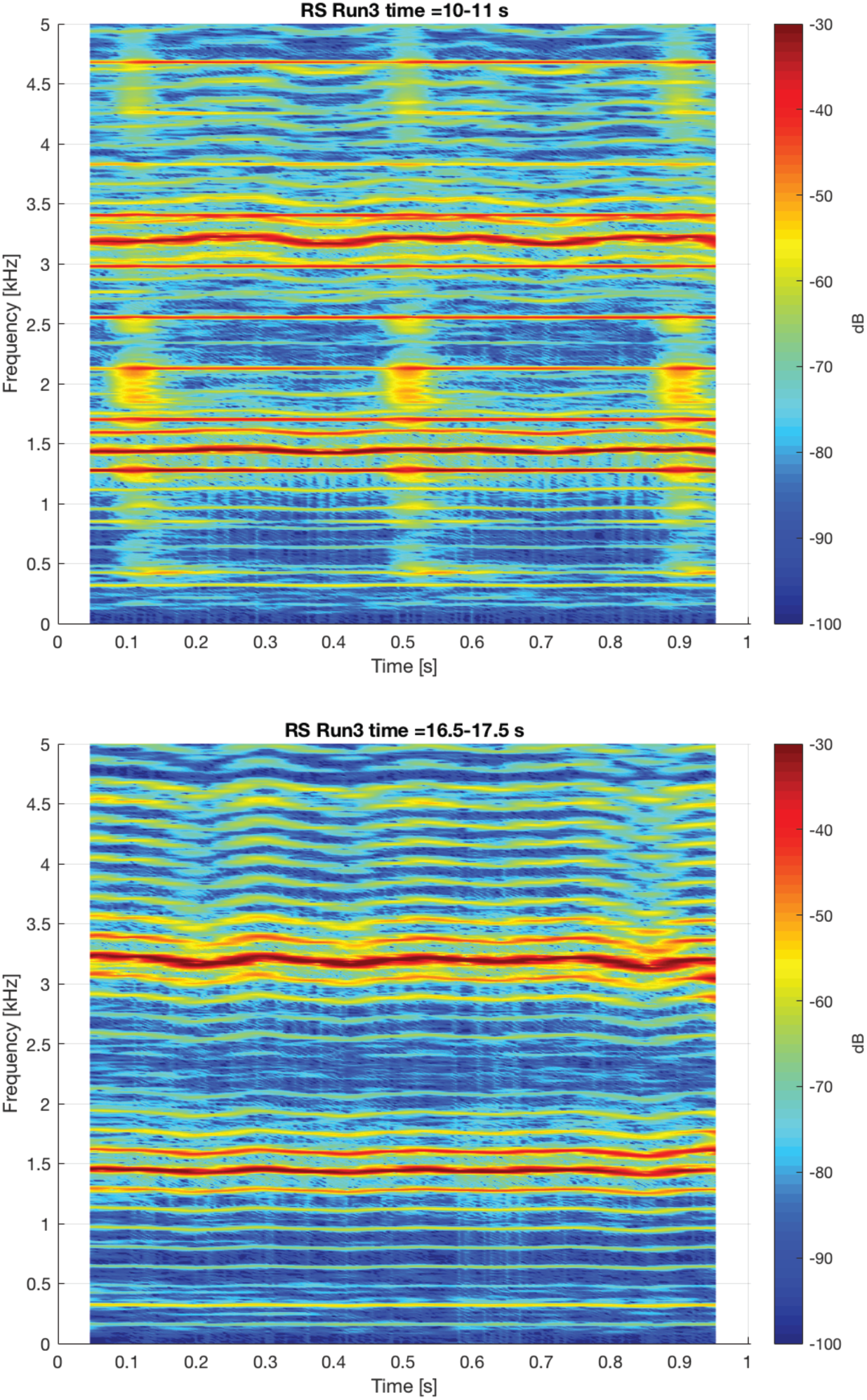
Spectrogram of steady-state overtone voicing whose volumetric scan is shown in Fig.19. Two different one-second segments are shown: top shows during the scan (and thus includes acoustic noise from the scanner during image acquisition), while bottom shows just after scan ends but subject continues to sing.

#### Vocal tract shape & Associated spectrograms

Examples of the vocal tract taken during the dynamic MRI runs (i.e., midsagittal only) are shown for a very different representative time points in Fig. 21.

**Appendix 1 Figure 21.**
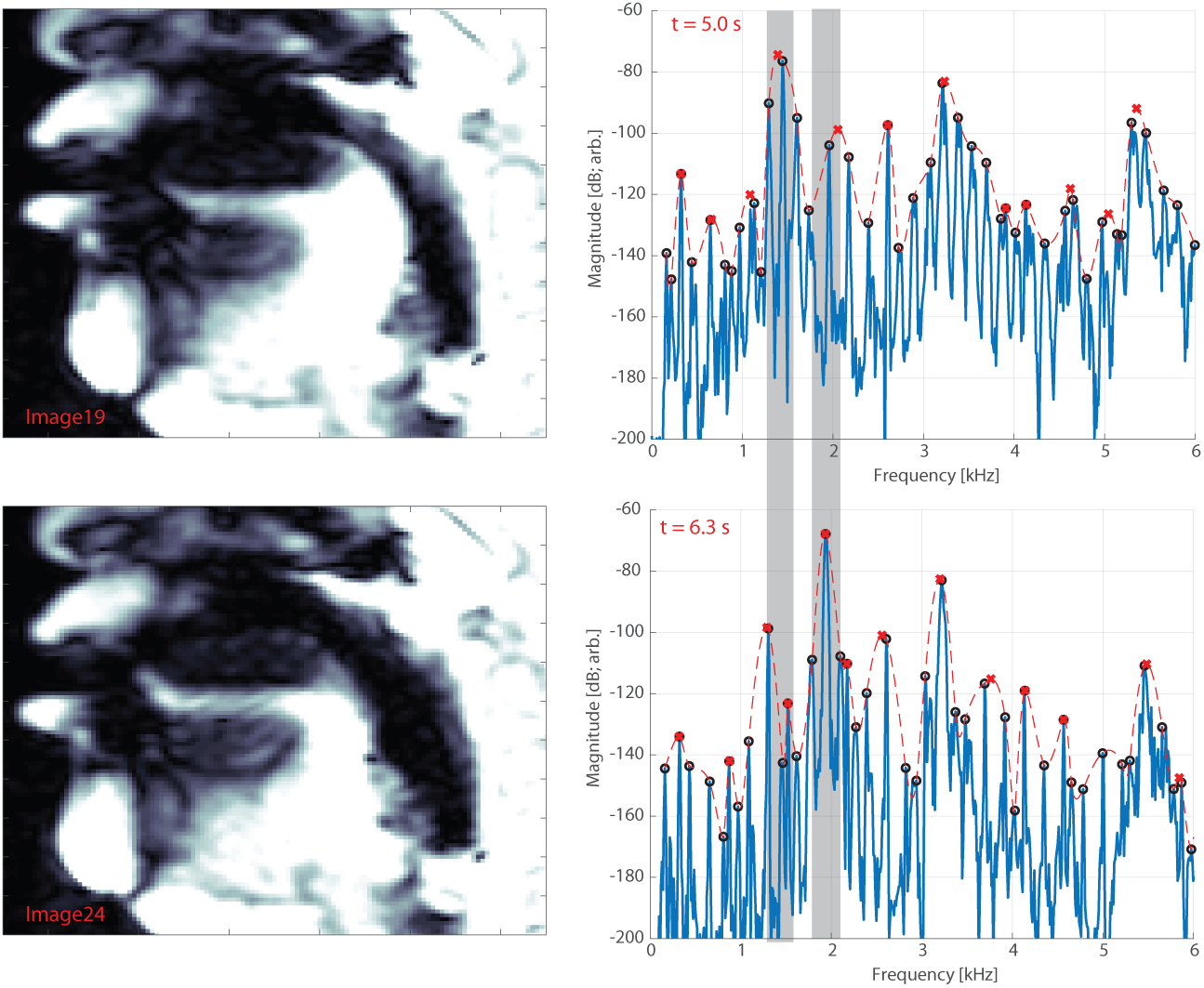
Representative movie frames and their corresponding spectra for singer T2, as input into modeling parameters (e.g., Fig.5). Corresponding *Appendix* data files are DynamicRun2S.mov (MRI images) and DynamicRun2sound.wav (spectra; see also DynamicRun2SGrid.pdf). Top row shows a “low pitch” (first) focused state at about 1.3 kHz while the bottom row shows a “high” pitch at approximately 1.9 kHz. Note a key change is that the back of the tongue moves forward to shift from the low to high pitch. Thin grey bars are added to the spectra to help highlight the frequency difference. Same legend as that shown in Fig.1.

### Acoustic data

All acoustic recordings can be accessed via the following links^*b*^:

- T1_1.wav – https://www.dropbox.com/s/ezwazkw3566fvdj/T1_1.wav?dl=0
- T1_2.wav – https://www.dropbox.com/s/kzixom608ea13ig/T1_2.wav?dl=0
- T1_3.wav – https://www.dropbox.com/s/2yr1mom32kjyqze/T1_3.wav?dl=0
- T1_3short.wav – https://www.dropbox.com/s/2h6z7qbauzj4yqa/T1_3short.wav?dl=0
- T2_1.wav – https://www.dropbox.com/s/srz9heab92ax0c2/T2_1.wav?dl=0
- T2_1shortA.wav – https://www.dropbox.com/s/r44cf8co149l5gi/T2_1shortA.wav?dl=0
- T2_1shortB.wav – https://www.dropbox.com/s/65hk1p5iklrdv0g/T2_1shortB.wav?dl=0
- T2_1shortC.wav – https://www.dropbox.com/s/njw3x41eced1co9/T2_1shortC.wav?dl=0
- T2_2.wav – https://www.dropbox.com/s/s9v6q7w4vgglnvt/T2_2.wav?dl=0
- T2_2short.wav – https://www.dropbox.com/s/azi533et1bv8xnk/T2_2short.wav?dl=0
- T2_3.wav – https://www.dropbox.com/s/vh0dfe7wyeafns4/T2_3.wav?dl=0
- T2_4.wav – https://www.dropbox.com/s/dilioeeoxmkzkli/T2_4.wav?dl=0
- T2_5.wav – https://www.dropbox.com/s/4cxeajghyzlsl0k/T2_5.wav?dl=0
- T2_5longer.wav – https://www.dropbox.com/s/36ky51tpzpmbevo/T2_5longer.wav?dl=0
- T2_5short.wav – https://www.dropbox.com/s/vq035otu8hp193m/T2_5short.wav?dl=0
- T3_2.wav – https://www.dropbox.com/s/nt07x0pux8ydtnu/T3_2.wav?dl=0
- T3_2shortA.wav – https://www.dropbox.com/s/9lzcsdr64nmani4/T3_2shortA.wav?dl=0
- T3_2shortB.wav – https://www.dropbox.com/s/iambc5myzrvedvk/T3_2shortB.wav?dl=0
- T4_1.wav – https://www.dropbox.com/s/e4pzrofunvo12wn/T4_1.wav?dl=0
- T4_1shortA.wav – https://www.dropbox.com/s/dwpihcag2zlu5jy/T4_1shortA.wav?dl=0

### MRI data

#### Images

Images were only obtained from singer T2. Files can be accessed via the following links. Note that all image data are saved as DICOM files (i.e., .dcm).:

- Volumetric Run1 – https://www.dropbox.com/sh/xsm4sjpdyyv81yh/AABlLIGw6GHn4T1vI4fXhwsIdl=0
- Volumetric Run2 – https://www.dropbox.com/sh/il8tji4jifl0fir/AADUAzPlviNqa-Q1JVHAzcx_a?dl=0
- Volumetric Run3 – https://www.dropbox.com/sh/64fuvhc51g9iik2/AADHYox_G65pBKDGgFGDwNdl=0
- Dynamic midsagittal Run1 – https://www.dropbox.com/sh/ca3foed5d02zs2a/AAALQATMKhMt8Qdl=0
- Dynamic midsagittal Run2 – https://www.dropbox.com/sh/oaqg2l12vl3fkn0/AAC16F1zhOQslC85dl=0
- Dynamic midsagittal Run3 – https://www.dropbox.com/sh/ts42h9r4plc6ghh/AAAne9VkcZ9OFhledl=0

#### Audio Recordings

Acquired during MRI acquisition (see Methods). Files can be accessed via the following links:

- Vol. Run1 audio – https://www.dropbox.com/s/q6y3ei1dgrfe2ln/Run1Vsound.wav?dl=0
- Vol. Run2 audio – https://www.dropbox.com/s/xp1hyly2y80q896/Run2Vsound.wav?dl=0
- Vol. Run3 audio – https://www.dropbox.com/s/cx1lps2nespew4c/Run3Vsound.wav?dl=0
- Dyn. Run1 audio – https://www.dropbox.com/s/f7dz08bq0dlhjiu/Run1sound.wav?dl=0
- Dyn. Run2 audio – https://www.dropbox.com/s/lzhfyooyjap29yz/Run2sound.wav?dl=0
- Dyn. Run3 audio – https://www.dropbox.com/s/wegw6j1v33jnn2v/Run3sound.wav?dl=0

#### MRI Movies

Midsagittal movies with sound were also created by animating the frames in Matlab and syncing the recorded audio via Wondershare Filmora. They are saved as .mov files (Apple QuickTime Movie file) and can be accessed via:

- Dyn. Run1 video – https://www.dropbox.com/s/kkoo7clxka2rrc0/DynamicRun1S.mov?dl=0
- Dyn. Run2 video – https://www.dropbox.com/s/lfnvjqrmj1mbd54/DynamicRun2S.mov?dl=0
- Dyn. Run3 video – https://www.dropbox.com/s/sguz1rrpps0odc1/DynamicRun3S.mov?dl=0

To facilitate connecting movie frames back to the associated sound produced by singer T2 at that moment, the movies include frame numbers. Those have been labeled on the corresponding time location in the spectrograms (see red labels at top), accessible here:

- Dyn. Run1 spectrogram – https://www.dropbox.com/s/nqfdm6xnxrgthh1/DynamicRun1SGrid.pdf?dl=0
- Dyn. Run2 spectrogram – https://www.dropbox.com/s/4wvc7hs96h8nel3/DynamicRun2SGrid.pdf?dl=0
- Dyn. Run3 spectrogram – https://www.dropbox.com/s/vpy8n3g0vc3rdoy/DynamicRun3SGrid.pdf?dl=0

#### Segmented volumetric data

Files (like those shown in Fig.3) can be accessed via the following links. Note that all data are saved as STL files (i.e., .stl).:

- Segmented data (T2) – https://www.dropbox.com/s/5fpxkkztvn0xntv/RTfullReconSmoothed.stl?dl=0

### Software & Synthesized song

#### Analysis software

Waveform analysis was implemented in Matlab and may be accessed via:

- Code to analyze general aspects of the waveforms (e.g., Fig.1 spectrograms) – https://www.dropbox.com/s/cotkuo5453u5ki0/EXspectrogramT6SI.m?dl=0
- Code to quantify *e*_*R*_ time course (e.g., Fig.2)– https://www.dropbox.com/s/uqtkbdw3lr93gxe/EXquantifyFocusedSI.m?dl=0

## Footnotes

1 *Pressed* phonation, also referred to as ventricular voice, occurs when glottal flow is affected by virtue of tightening the laryngeal muscles such that the ventricular folds are brought into vibration. Such has the perceptual effect of adding a degree of roughness to the voice sound. An anecdotal popular fiction account along these lines appears in the 2005 American film The Exorcism of Emily Rose.

2 It should be noted that ***Herzel and Reuter (1996)*** go so far to define *biphonation* explicitly through the lens of nonlinearity. We relax such a definition and argue for a perceptual basis for delineating the boundaries of biphonation.

a Citations here presently refer to the bibliography of the main document. Also, because sections in the Appendix are not enumerated (due to the template), the section naming here is a bit clunky.

b These are currently active Dropbox links. However upon publication, these links would change to the journal’s online repository

